# Spike-frequency dependent coregulation of multiple ionic conductances in fast-spiking cells forces a metabolic tradeoff

**DOI:** 10.1101/2021.03.08.434486

**Authors:** Yue Ban, Rosalie Maltby, Michael R. Markham

**Affiliations:** Department of Biology, University of Oklahoma, Norman, OK 73019, USA; Cellular & Behavioral Neurobiology Graduate Program, University of Oklahoma, Norman, OK 73019, USA

## Abstract

High-frequency action potentials (APs) allow rapid information acquisition and processing in neural systems, but create biophysical and metabolic challenges for excitable cells. The electric fish *Eigenmannia virescens* images its world and communicates with high-frequency (200-600 Hz) electric organ discharges (EODs) produced by synchronized APs generated at the same frequency in the electric organ cells (electrocytes). We cloned three previously unidentified Na^+^-activated K^+^ channel isoforms from electroctyes (eSlack1, eSlack2, and eSlick1). In electrocytes, mRNA transcript levels of the rapidly-activating eSlick, but not the slower eSlack1 or eSlack2, correlated with EOD frequency across individuals. In addition, transcript levels of an inward-rectifier K^+^ channel, a voltage-gated Na^+^ channel, and Na^+^,K^+^-ATPases also correlated with EOD frequency while a second Na^+^ channel isoform did not. Computational simulations showed that maintaining electrocyte AP waveform integrity as firing rates increase requires scaling conductances in accordance with these mRNA expression correlations, causing AP metabolic costs to increase exponentially.

## INTRODUCTION

Organisms expend precious metabolic energy to acquire, process, and store information. Action potentials (APs) are central to these processes because they are a fundamental unit of information in nervous systems. As a result, the rules that govern these energy-information tradeoffs are often revealed in the biophysical mechanisms of AP generation [1, 2]. APs in excitable cells are initiated by depolarizing inward Na^+^ currents and terminated by repolarizing outward K^+^ currents. Each AP incurs a metabolic cost when the Na^+^,K^+^-ATPases hydrolyze ATP to restore the transmembrane Na^+^ and K^+^ gradients after each AP. The kinetics and densities of the Na^+^ and K^+^ ion channels that generate these APs determine the waveform of each individual AP, the maximum AP firing rate and, ultimately, the metabolic costs of AP generation. Sustaining high AP firing frequencies supports rapid information acquisition and processing [3–5], but presents a significant metabolic challenge for two reasons. First, the very feature of high firing rates imposes metabolic costs by virtue of more APs per unit time. Second, maintaining high firing rates requires very brief APs, and the generation of brief APs often requires the metabolically inefficient overlap of depolarizing Na^+^ currents and repolarizing K^+^ currents [6].

The freshwater weakly electric fish *Eigenmannia virescens* generates high-frequency electric organ discharges (EODs) to image their world and communicate in darkness (Fig. 1) [7]. *E. virescens* has a considerable animal-to-animal variability in EOD frequency (EODf) which is set by a medullary pacemaker nucleus, with each fish maintaining a relatively fixed frequency in the range of 200-600 Hz [8]. Early studies on the behavior of *E. virescens* suggested that EODf conveys information about individual identity, gender and dominance rank [7, 9]. These high-frequency EODs confer two major advantages by providing fast sensory sampling rates and shifting the signal energy away from low frequencies that are detectable by electroreceptive predators [10], but high-frequency EODs also incur staggering metabolic costs [11].

**Fig. 1.**
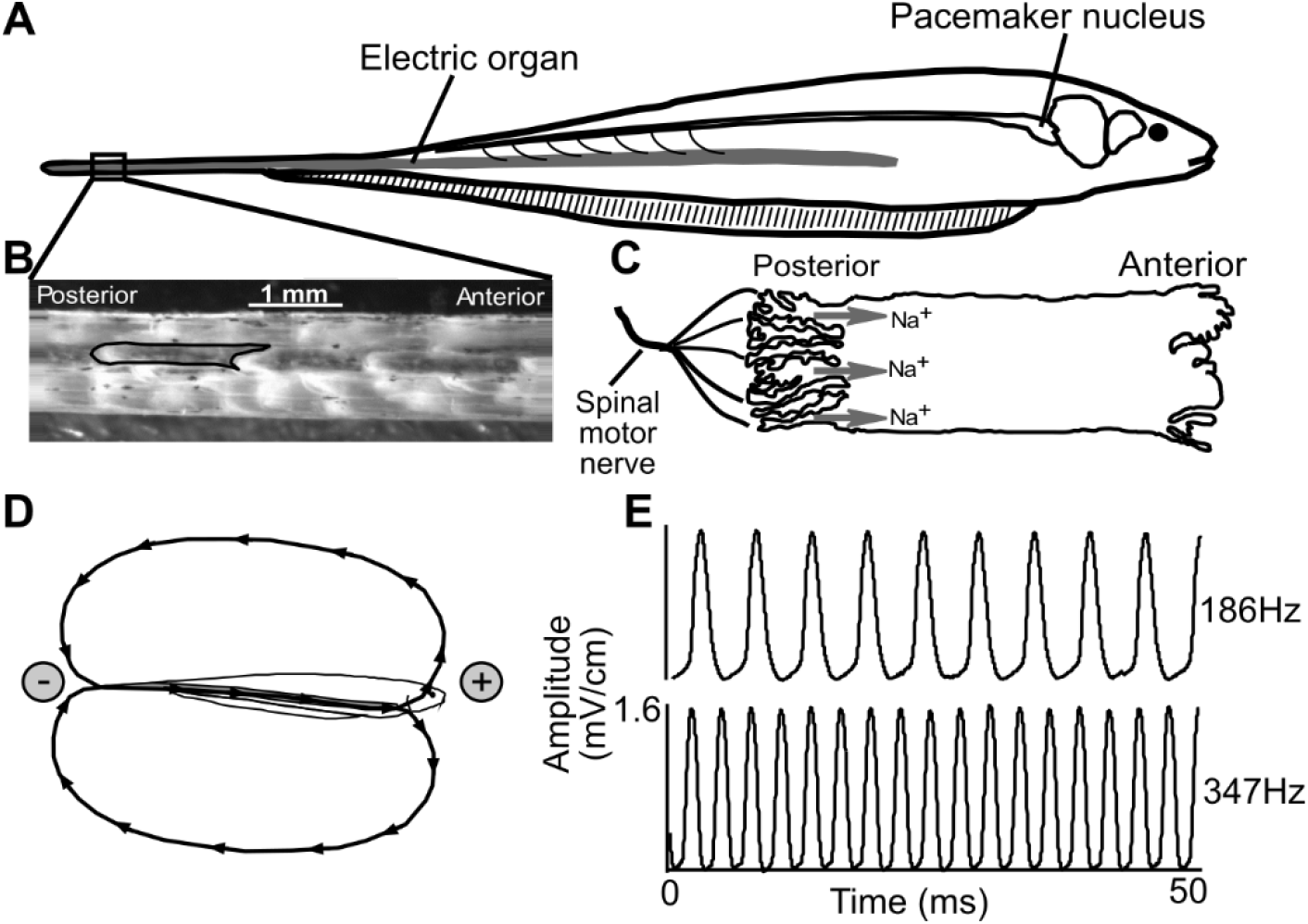
EOD generation in *E. virescens*. A: The EO runs longitudinally along the fish body and extends into the caudal tail filament. *B:* A section from the tail with skin removed to expose the EO. A single electrocyte is outlined in black. *C:* Schematic of an electrocyte. Electrocytes are highly polarized cells approximately 1.5 mm in anterior-posterior length and 0.6 mm in diameter. Electrocyte APs are controlled by the medullary pacemaker nucleus via spinal motor neurons innervating on the posterior membrane of each electrocyte. The cell’s innervated posterior face is deeply invaginated and occupied by cholinergic receptors and voltage gated Na^+^ (Na_v_) channels. The activation of cholinergic synapses causes an inward Na^+^ current. *D:* The Na^+^ current moves toward the head, and followed by a return path from head to tail in the surrounding water. *E:* The EOD waveforms recorded from fish with high and low EOD frequency. Panels A-D were modified from Ban et al., 2015.

The EOD is produced by more than 1000 electric organ cells (electrocytes) that generate near simultaneous APs at the EOD frequency (200-600 Hz), approximately the same frequency range as the maximum sustained firing rates for fast-spiking mammalian cortical neurons [12] and brainstem auditory neurons [13]. However, electrocytes maintain these high frequencies unremittingly throughout the lifespan, whereas central neurons sustain such high firing rates only for brief periods. Further, in addition to the biophysical and metabolic challenges associated with maintaining constant high-frequency APs, the resulting metabolic costs are amplified in *E. virescens* electrocytes because electrocyte ionic currents can exceed 10 μA during each AP, with corresponding entry of approximately 2 x 10^10^ Na^+^ ions into the cell during each AP [11, 14]. These large ionic currents create extreme metabolic demands because one ATP must be hydrolyzed by the Na^+^,K^+^-ATPase for every three Na^+^ ions returned to the extracellular space. High firing rates also create extreme demand for rapid charge translocation by the Na^+^,K^+^-ATPases because the interval for restoring ionic gradients between APs ranges from just 2.5 ms to as little as 0.8 ms across the EODf range of 200-600 Hz.

Here we investigated the ionic mechanisms associated with high-frequency APs in *E. virescens* electrocytes and the associated demands on Na^+^,K^+^-ATPase activity. Electrocytes are highly polarized cells approximately 1.5 mm in length and 500 μm in diameter. APs are initiated with the activation of cholinergic receptors and Na_v_ channels on the innervated posterior membrane to allow the influx of Na^+^ (Fig. 1). Electrocytes in *E. virescens* terminate APs with Na^+^-activated K^+^ (K_Na_) channels rather than the voltage gated K^+^ (Kv) channels that terminate the AP in the electrocytes of related species [14–17].

Numerous studies have suggested that the density and kinetic properties of K^+^ channels in the plasma membrane are key determinants of an excitable cell’s functional capacity [18–23]. Therefore, we cloned the cDNAs encoding the K_Na_ channels in *E. virescens* EOs and identified three different types of K_Na_ channel subunits expressed in electrocytes. Two of these channels, eSlack1 and eSlack2, closely resemble K_Na_ channels encoded by the Slack gene in mammalian systems; and the third channel, eSlick, shares the highest homology to the Slick channel in rat. By expressing fluorescent protein tagged K_Na_ channel subunits in electrocytes, we showed that all three K_Na_ channels are expressed on the cell’s anterior region, separated by >1 mm from the Na_v_ channels which are restricted to the posterior membrane.

We also examined the functional differences among the three K_Na_ channels by expressing them in *X. laevis* oocytes. Recordings of whole cell currents showed that eSlick currents are activated much more rapidly than eSlack1 currents. To explore which conductances play key roles in determining the firing frequency of electrocytes, we used qRT-PCR to measure the mRNA levels of genes encoding ion channels and Na^+^/K^+^ ATPases in EO from fish with different EODf. The transcription levels of eSlick, Nav1.4a, Kir6.2 and Na^+^/K^+^ ATPase increase with EODf, while transcription levels of eSlack1, eSlack2, Nav1.4b did not correlate with EODf.

In computational simulations of electrocytes stimulated at a broad range of EODfs, we found that maintaining AP integrity across firing rates required scaling ionic conductances in accordance with our experimentally derived mRNA expression correlations. These simulations also revealed that AP metabolic costs and the rate of required charge translocation by the Na^+^,K^+^-ATPases increased exponentially with higher frequencies.

## RESULTS

### Molecular identities of K_Na_ channels in *E. virescens* electrocytes

Mammalian K_Na_ channels are encoded by two highly similar paralog genes, *Slo2.1 (Slick, kcnt2*) and *Slo2.2*(*Slack, kcnt1*) belonging to the *Slo* gene family [24, 25]. In *E. virescens* EOs, we cloned three full-length cDNAs similar to the mammalian *Slo2* transcripts. Phylogenetic analysis (Fig. 2A) of channels in the SLO family show that two cloned cDNAs have the strongest homology with mammalian *Slack* transcripts, and the third full-length cDNA is more closely related to known *Slick* transcripts. The open reading frames (ORFs) of the two *E. virescens* Slack genes encode two proteins that consist of 1164 and 1030 amino acids, respectively, which share 68.6% homology. Amino acid differences between the two *Slack* proteins were dispersed along the entire sequence, suggesting they are not likely generated by RNA alternative splicing. Given the evidence that duplication of voltage gated sodium and potassium channel genes has occurred in multiple gymnotiform species [21, 26], gene duplication is more likely the mechanism giving rise to the two Slack transcript variants in *E. virescens*. We designated the duplicated *Slack* genes in *E. virescens* as *eSlack1* (ORF: 3495 nt) and *eSlack2* (ORF: 3093 nt). Therefore, *E. virescens* EOs express three different K_Na_ channel subunits, encoded by both *Slack* and *Slick* genes.

**Fig. 2.**
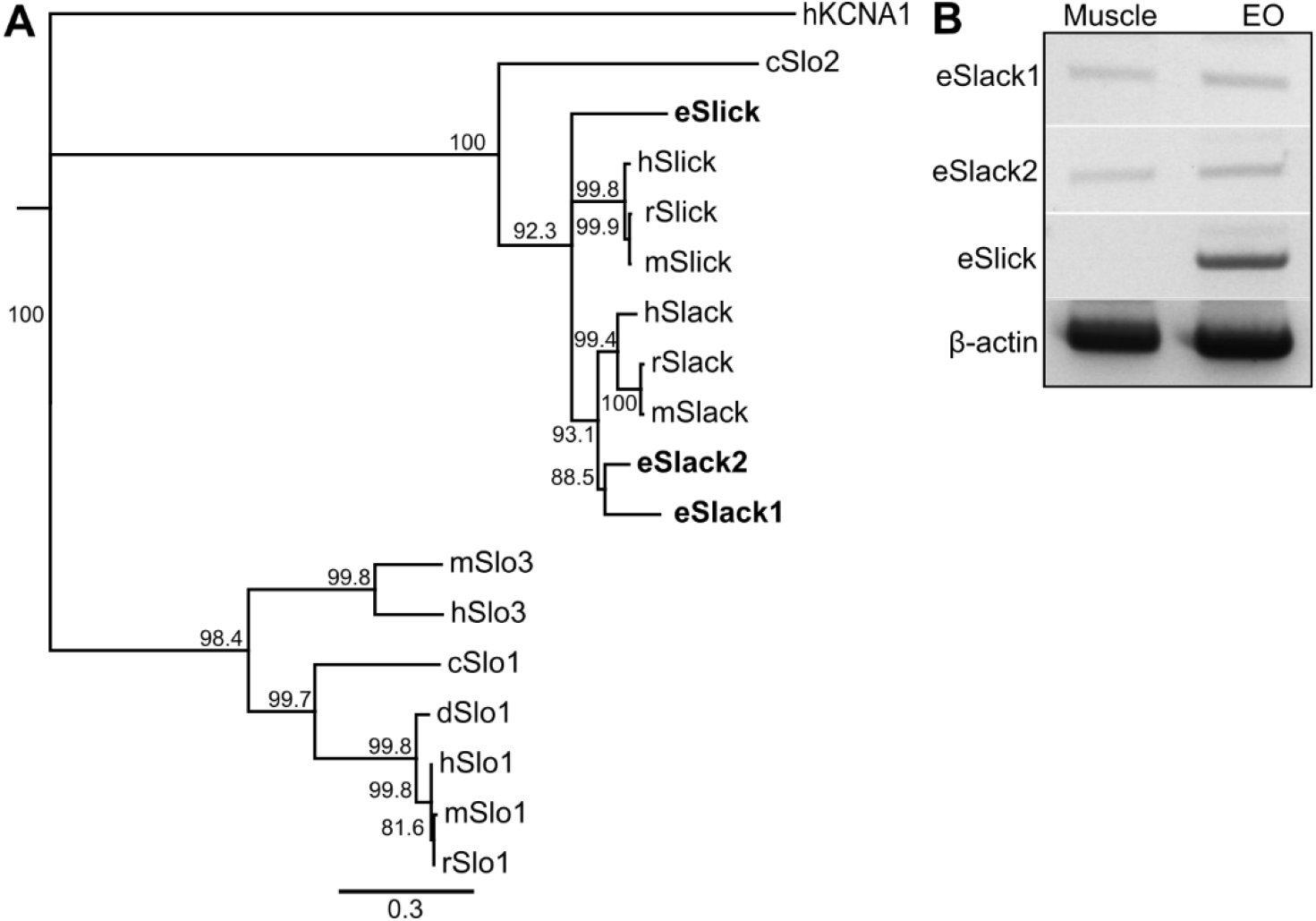
Molecular identities of *E. virescens* K_Na_ channel genes. *A:* A rooted neighbor-joining phylogenetic tree for the high conductance potassium channels in the SLO family. The family of SLO channels includes Slo1 (the “big” potassium (BK) K_Ca_ channel), Slo2.1 (the Slick K_Na_ channel), Slo2.2 (the Slack K_Na_ channel), and Slo3 (the large conductance pH-sensitive K^+^ channel). Human Kv1.1 was used as the outgroup. (h: *Homo sapiens;* r: *Rattus norvegicus;* m: *Mus musculus;* d: *Danio rerio;* c: *Caenorhabditis elegans;* e: *Eigenmannia virescens*). *B:* Expression pattern of *E. virescens* K_Na_ channels in muscle and EO. eSlack1 and eSlack2 were amplified from the cDNA of both muscle and EO, whereas eSlick was only amplified from EO cDNA. Primers and amplicon sizes are listed in Table 1.

In all but one genus of weakly electric fish, electric organs (EOs) are derived from skeletal muscle [27]. With the development and maturation of EOs, electrocytes eliminate the coupling between contraction and excitability [28, 29]. Due to the myogenic origin of EO tissue, we examined the expression pattern of eSlack1, eSlack2, and eSlick in *E. virescens* muscle and EO by reverse transcription PCR and found that eSlack1 and eSlack2 are expressed in both EO and muscle, whereas Slick is expressed only in the EO (Fig. 2B).

### Sequence and Structure of *E. virescens* K_Na_ Channels

The *E. virescens* eSlack1 and eSlack2 channel subunits share 74.3% and 70.8% homology to rat Slack-A, respectively [30]. Consistent with the structure of mammalian Slack channel subunits, both eSlack1 and eSlack2 subunits are predicted to contain six membrane-spanning domains (S1-S6) with a pore-forming loop between S5 and S6, and an extensive cytoplasmic C-terminal region (Fig. 3) [24, 31–33]. Slack channels are activated by intracellular Na^+^ ions, and the sensitivity of these channels to Na^+^ is determined by the presence of a Na^+^ coordination motif in the second Regulator of K^+^ conduction (RCK) domain. This motif contains six amino acids in rat Slack subunits (<u>D</u>NKPD<u>H</u>), with aspartic acid (D) and histidine (H) in the beginning and ending position [34]. In the homologous position, *E. virescens* Slack-1 and Slack-2 subunits have the sequence <u>D</u>NQPDD<u>H</u> and <u>D</u>NPPDN<u>H</u> respectively, making them putative Na^+^ binding sites (Fig. 3A). There is great divergence between *E. virescens* Slack-1 and Slack-2 in the N-and C-terminus. eSlack2 has a C-terminal tail approximately 100 amino acids shorter than eSlack1 and all other identified Slack and Slick subunits in mammals (Fig. 2A). The N-terminus is where amino acid differences are most frequently found between eSlack1 and eSlack2. The N-terminus of eSlack2 is highly similar to that of mouse and rat Slack-A. In mammals, RNA alternative splicing gives rise to three Slack transcripts, Slack-A, Slack-B and Slack-M, which are regulated by alternative promoters and differ in the N-terminal residues [30]. eSlack1 has a unique N-terminus which is not similar to any known mammalian Slack isoforms (Fig. 3).

**Fig. 3.**
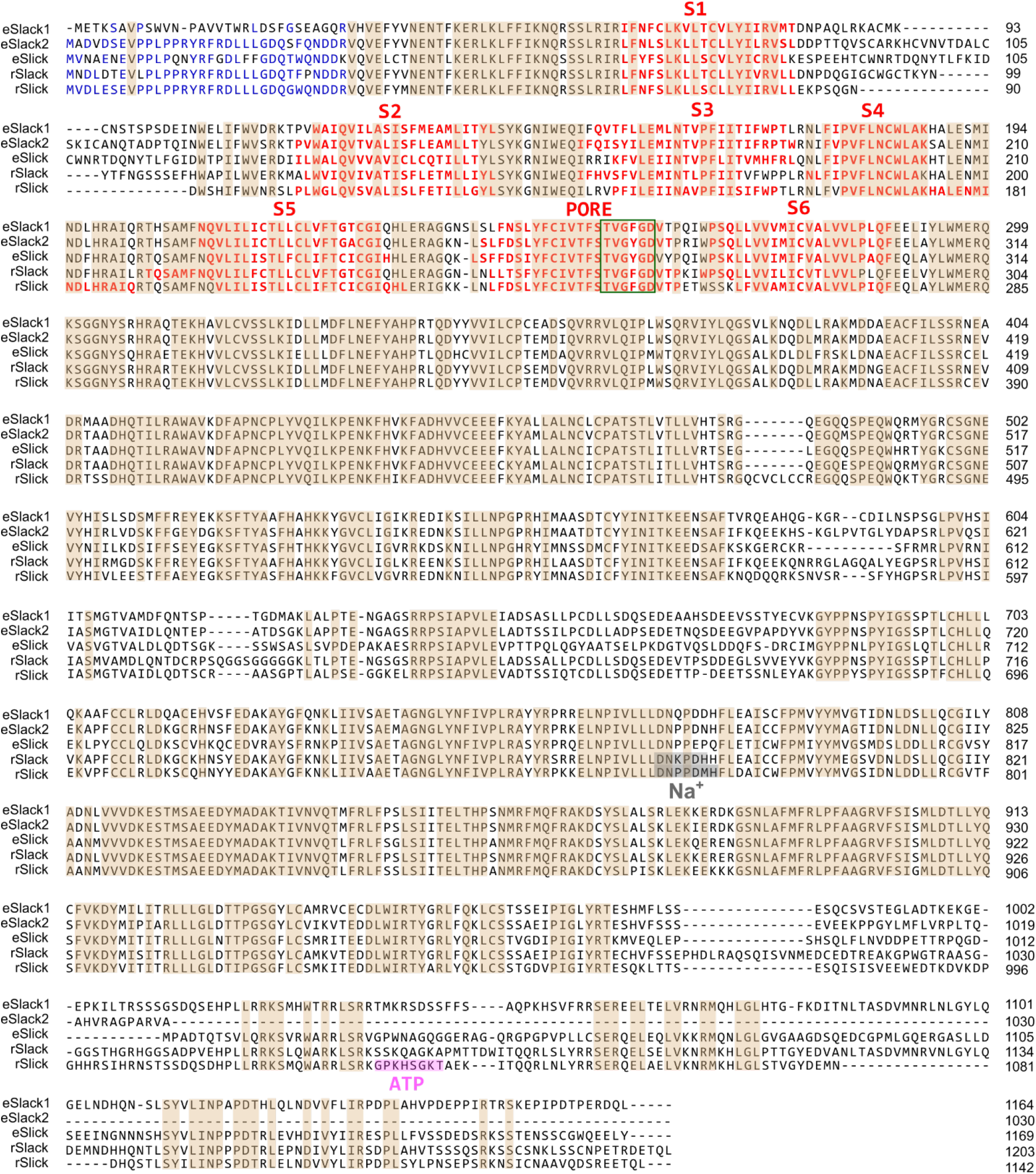
Amino acid sequences of *E. virescens* K_Na_ channels. *A:* Multiple sequence alignment was run by the Clustal W program using the *Geneious* software. Identical amino acids among all five sequences are shaded in brown except the C-terminal tail, where residues shared by eSlack1, eSlick, rSlack and eSlick are also highlighted. Gaps are represented by dashed lines. In the cytoplasmic N-terminus, identical residues are colored in blue. Red residues represent the membrane spanning domains (S1-6) and the pore region (P) of the five K_Na_ channels. Within the pore forming loop, the conserved residues determining the channel’s specific selectivity to K^+^ ions are highlighted with green box. Residues shaded in gray represent the Na^+^ coordination motifs in rat Slack and Slick. Residues composing the ATP binding motif of rat Slick are shaded in magenta.

The ORF of *E. virescens* Slick encodes a protein composed of 1142 amino acids, sharing 66.6% homology with rat Slick subunits. It has an N-terminus closely resembling that of eSlack2 and rat Slick, six predicted membrane spanning domains (S1-S6) with a pore-forming loop between S5 and S6, and an extensive C-terminal region (Figs. 3,4). At the homologous position of the Na^+^ coordination motif of rat Slick subunit (<u>D</u>NPPDM<u>H</u>) [35], *E. virescens* Slick has the sequence <u>D</u>NPPEPQ, which shares four of seven residues in common with rat Slick, and does not end with histidine (H). Histidine (H) may not be necessary for binding Na^+^, as it was shown in rat Slick that mutation of the aspartic acid (D) residue dramatically decreased the channel’s sensitivity to Na^+^, whereas histidine (H) substitution barely changed the channel’s function [35]. Rat and human Slick channels are ATP-regulated channels, and can be directly inhibited by intracellular ATP. The molecular determinants of ATP sensitivity is the presence of the “Walker A motif” (GxxxxGKT) on the distal C-terminus of Slick subunits [25, 36]. The residues at the homologous position of the “Walker A motif” in rat Slick are not well conserved between rat and *E. virescens* Slick. Furthermore, there is no motif having the signature residues of the “Walker A motif” in the C-terminus of *E. virescens* Slick subunits. Whether *E. virescens* Slick channels are regulated by intracellular ATP levels needs to be determined by future electrophysiology studies.

**Fig. 4.**
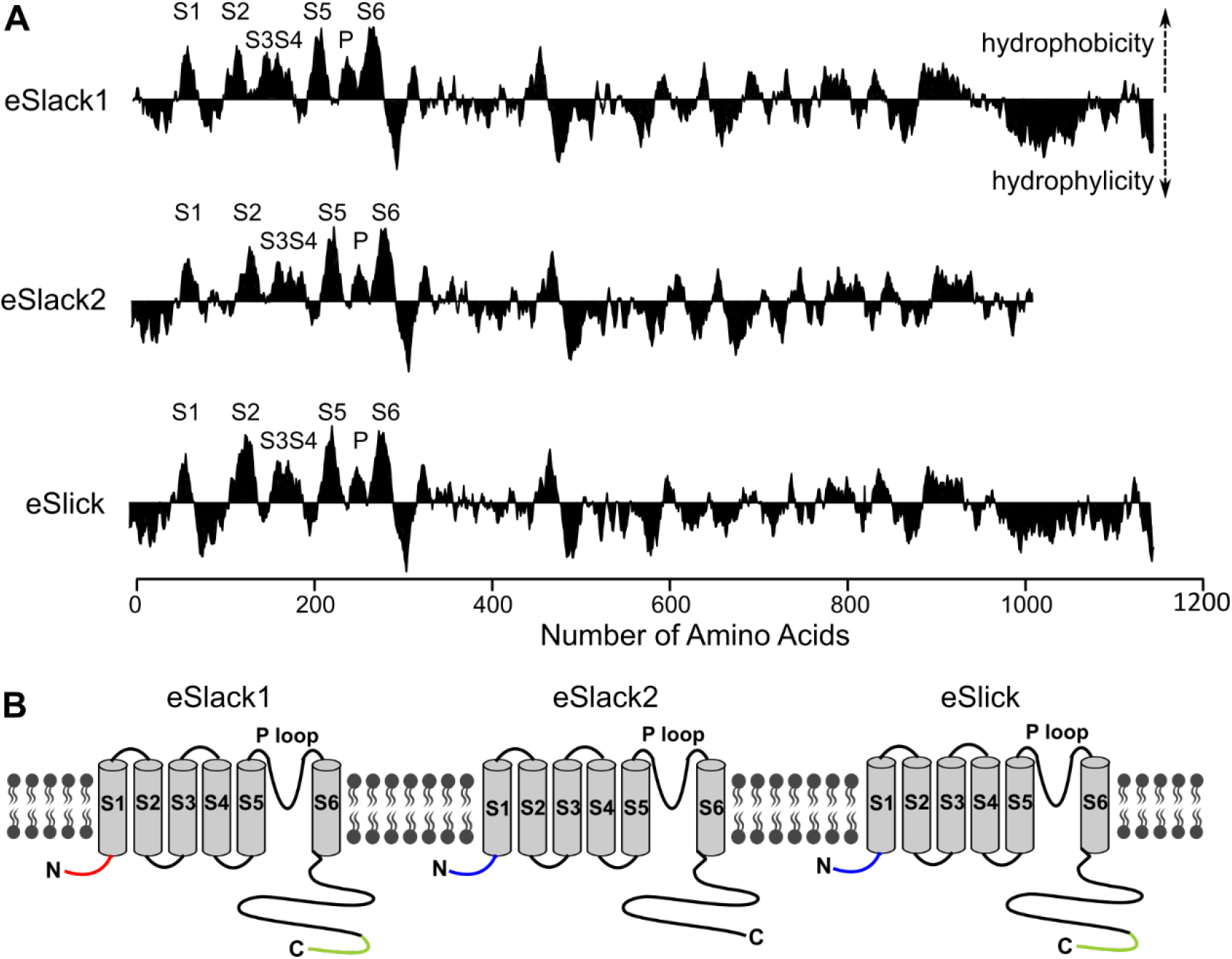
Predicted secondary structure of *E. virescens* K_Na_ channels. *B:* Kyte-Doolittle hydrophilicity plot of *E. virescens* K_Na_ channels (window size of 19 amino acids). *C:* Schematic representation of *E. virescens* K_Na_ channel subunits. eSlack2 and eSlick have identical N-terminus (blue), which is different from that of eSlack1(red). The C-terminal tail of eSlack2 is shorter than that of eSlack1 and eSlick (green).

### Expression patterns of Slack and Slick channels in electrocytes

The subcellular localization of ion channels plays a key role in determining the bioelectrical properties of excitable cells, especially for electrocytes, which are highly polarized cells with structurally and biophysically different posterior and anterior membranes. Our previous immunohistochemical studies revealed that K_Na_ channels are located on the anterior region, separated by >1 mm from cholinergic receptors and Na^+^ channels restricted to the posterior membrane [37]. Protein sequence alignment between the peptide immunogen of an anti-Slack antibody we previously validated with *E. virescens* (Aviva Systems Biology, OAAJ11822) [38] and the three *E. virescens* K_Na_ subunits showed that the K_Na_ channel antibody likely targeted only the eSlack1 subunit. Due to the lack of specific commercially-available antibodies and the failure of multiple custom-generated antibodies to produce specific labeling of eSlack2 and eSlick channels, we took the approach of expressing fluorescent protein tagged constructs of these ion channel subunits to visualize the location of eSlack2 and eSlick subunits in electrocytes. It has been shown that direct injection of naked DNA plasmids encoding transgenes produced expression of those transgenes in fish muscle [39]. Electrocytes of *E. virescens* can reliably express fluorescent protein tagged actin following bulk injection of Actin-GFP expression vectors into the EO (M. Markham and H. Zakon: unpublished observations). Because electrocytes exhibit autofluorescence with excitation and emission spectra similar to those of green fluorescent protein [37], we therefore used a red fluorescent protein (mCherry) as the tag to construct the recombinant *E. virescens* K_Na_ channel subunits.

mCherry was fused to the N-terminus of eSlack1, eSlack2,, and eSlick subunits and separated by a flexible polylinker containing glycine (G) polypeptide with serine (S) inserts to allow the proper folding and function of both molecules [40–42]. The fusion of a fluorescent protein to a native protein may affect the protein’s normal localization. To ensure N-terminal mCherry fusion does not affect the trafficking of eSlack/Slick subunits to the plasma membrane, we examined the membrane expression of these recombinant K_Na_ channel subunits in *Xenopus laevis* (*X. laevis*) oocytes and showed that all of them could be successfully expressed on cell membranes (Fig. 5A). Next we injected mCherry-eSlack1, mCherry-eSlack2, or mCherry-eSlick expression vectors into the EO and performed live-cell imaging of electrocytes 10 days later. We found that mCherry-eSlack1 localized on the anterior region of the electrocyte, mimicking the distribution of endogenous eSlack1 detected by immunohistochemistry [38] (Fig. 5C), providing evidence that N-terminal mCherry fusion does not affect the normal localization of eSlack/Slick subunits. Similar to mCherry-eSlack1, the expression of mCherry-eSlack2 and mCherry-eSlick was only detected on the anterior region (Fig. 5D,E). These results indicate that the three *E. virescens* K_Na_ channel subunits all are expressed only on the anterior region of the electrocyte.

**Fig. 5.**
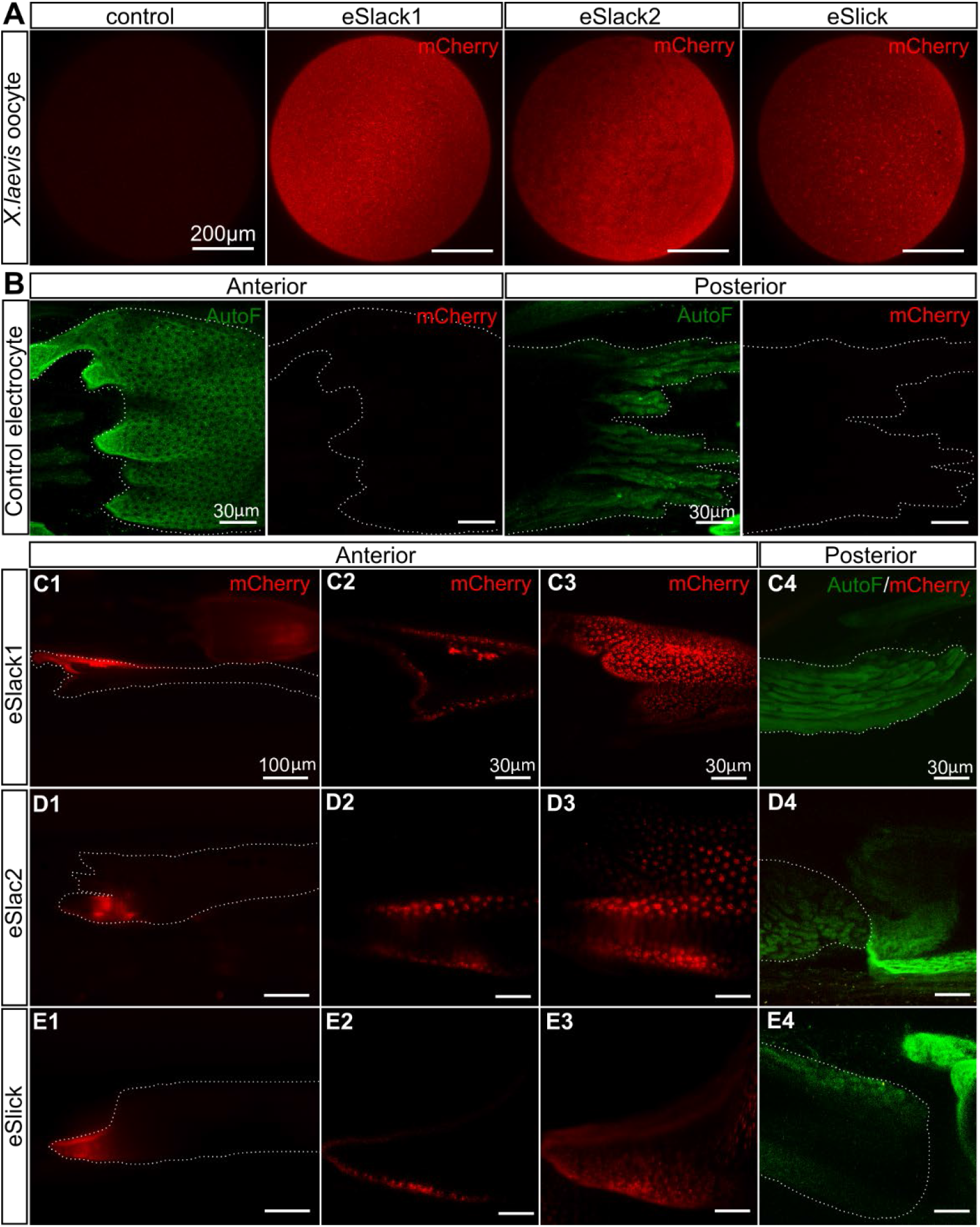
Expression of mCherry-tagged K_Na_ channels on the plasma membrane of *X. laevis* oocytes and localization of K_Na_ channels in electrocytes. A. Maximum-intensity-projection (MIP) images of *X. laevis* oocytes expressing mCherry tagged *E. virescens* K_Na_ channels rendered with images taken at different focal planes. B: MIP images of a control electrocyte (no expression of mCherry-tagged K_Na_ channels). Broadband tissue autofluorescence (AutoF, green) was excited by a 488-nm laser. *C-E:* Representative images of electrocytes expressing mCherry-eSlack/Slick plasmids on the anterior region. Images in C1, D1 and E1, acquired using an epifluorescent microscope, show a larger field of view. Other images in C-E were acquired by laser-scanning confocal microscope. Images displayed in C2, D2 and E2 are single optical sections showing the anterior face from cells expressing recombinant K_Na_ channels (red). C3, D3 and E3 are MIP images rendered from the serial optical sections shown in C2, D2 and E2. Merged images of autofluorescence (green) and mCherry (red) in C4, D4 and E4 revealed that recombinant K_Na_ channels are not expressed on the posterior membrane of electrocytes. White dotted lines indicate the boundary of electrocytes.

### Characteristics of eSlack/eSlick currents

The only outward K^+^ current in *E. virescens* electrocytes is a noninactivating Na^+^-activated K^+^ current (IK_Na_) [14]. To determine how the three K_Na_ channels identified here contribute to outward K^+^ currents, we expressed these K_Na_ channels in *X. laevis* oocytes to characterize and compare their electrophysiological properties. Both eSlack1 and eSlick constructs produced robust outward K^+^ currents. Whole-cell currents from cells injected with eSlack1 cRNA showed much slower activation than those injected with eSlick cRNA (Fig. 6A). At test potentials positive to −40 mV, eSlack1 activates with relatively slow time constants (Fig. 6A and B). The τ-V relationship eSlack1 activation shows that eSlack1 currents activate slower as membrane potential becomes more depolarized until +10 mV, after which activation becomes more rapid with more depolarized membrane potentials (Fig. 6C). In contrast, eSlick shows rapid activation. Whole-cell currents in eSlick cRNA injected oocytes activated nearly instantaneously with step changes in voltage to test potentials positive to −70mV (Fig. 6A and B). eSlick activation τ decreases with more depolarized membrane potentials reaching a minimum at +20 mV (Fig. 6D). The plateau is likely due to the inward-rectification. We used water-injected oocytes as controls. As reported previously, control cells could express an endogenous Ca^+^-activated Cl^-^ current and Na^+^-activated K^+^ current, the magnitude of which is much smaller compared to cells expressing exogenous currents (Fig. 6A) [43, 44]. Unlike for eSlack1 and eSlick, oocytes injected with eSlack2 cRNA expressed currents which are not distinguishable from those of control cells (Fig. 6A). The C-terminal tail of eSlack2 is approximately 100-amino acids shorter than eSlack1 and eSlick. Since mCherry-eSlack2 can be expressed on the plasma membrane of both *X. laevis* oocytes and electrocytes, the absence of currents is not likely due to the difficulty of trafficking eSlack2 into the plasma membrane. A likely possibility is that eSlack2 cannot form functional homotetrameric channels without the intact C-terminus.

**Fig. 6.**
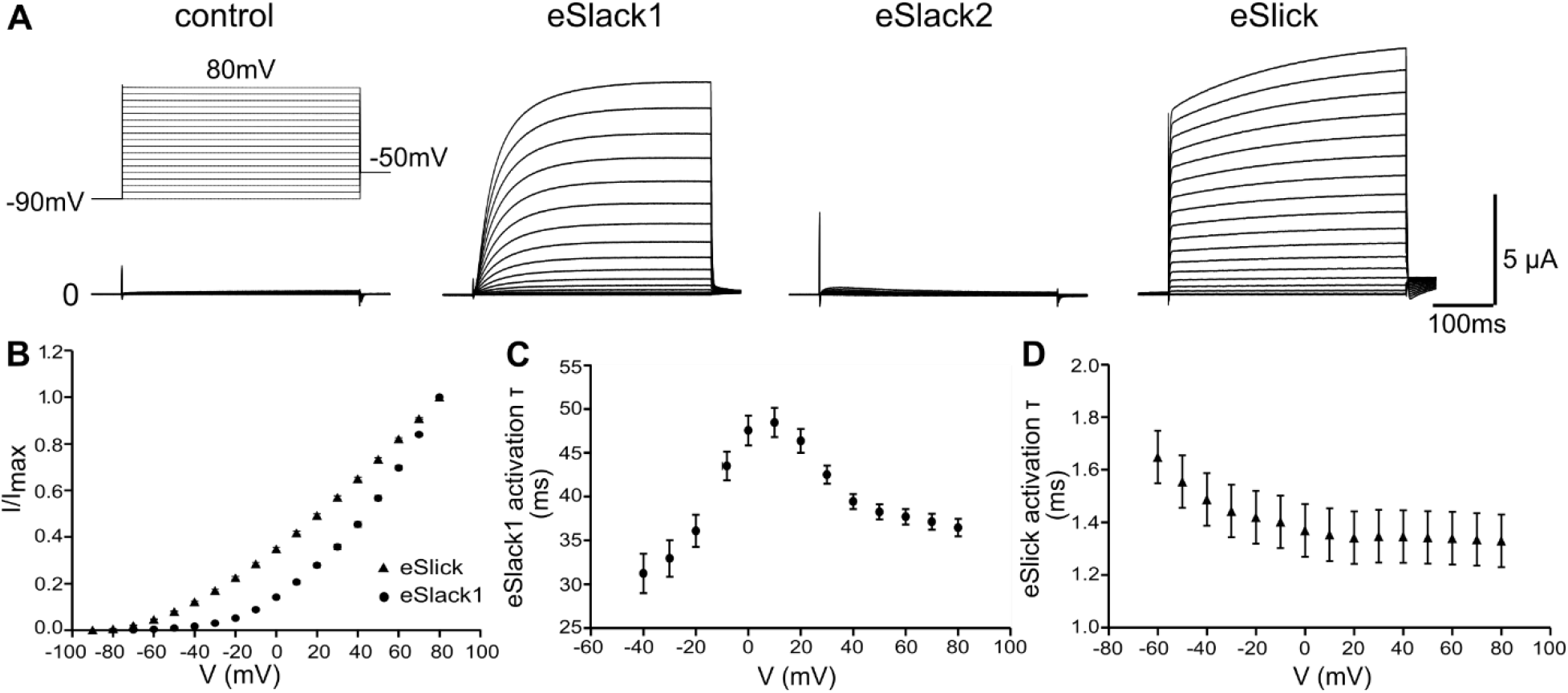
Whole cell recordings of *X. laevis* oocytes expressing *E. virescens* K_Na_ channels. *A:* Whole cell currents recorded when oocytes were depolarized by 400 ms voltage steps ranging from −90 mV to +80 mV in 10 mV increments every 5 s from a holding potential of −90 mV. *B:* Current-voltage relationship of oocytes expressing eSlack1 (circle; n=13) and eSlick (triangle; n=10) channels. Current amplitude was measured as the mean amplitude during the last 30 ms of each pulse, and divided by the maximal current amplitude. *C-D:* Activation time constant (τ) of eSlack1 (C; n=13) and eSlick (D; n=10) currents were plotted as a function of membrane potential.

The activity of mammalian Slack and Slick channels are regulated by the intracellular levels of Na^+^ and Cl^-^. We therefore examined the effects of elevated [Na^+^]_i_ on the activity of eSlack1 and eSlick channels. Thomson et al. showed that filling low resistance microelectrodes with 2M NaCl would allow Na^+^ to diffuse into the cell and thereby increase [Na^+^]_i_ [35, 45]. We applied the same method to increase [Na^+^]_i_ and measured the amplitude of currents when the cell was depolarized to +20 mV repetitively from −90 mV every 10 s. The peak amplitude of eSlack1 and eSlick currents was higher than that of control cells immediately after impaling the cell, suggesting that eSlack1 and eSlick have basal levels of activity even at intraoocyte Na^+^ concentrations of ~10 mM [46] (Fig. 7A).

**Fig. 7.**
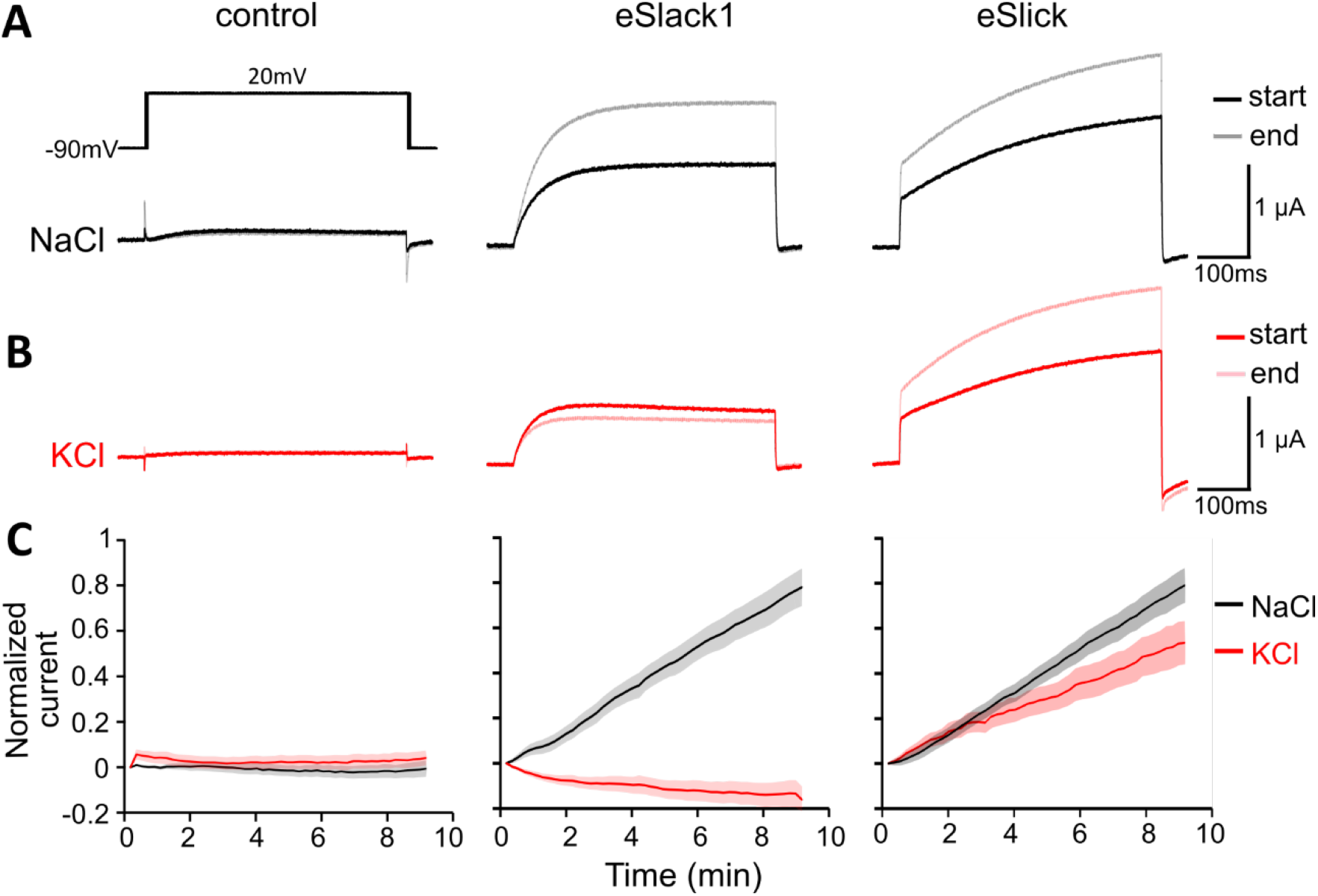
Whole cell currents ofeSlack1 and eSlick with microelectrodes filled with 2M NaCl or KCl. Oocytes were depolarized by a 500 ms pulse to +20 mV from a holding potential of −90 mV every 10s. *A:* With both microelectrodes filled with 2M NaCl, whole cell currents recorded from control oocytes (left), oocytes expressing eSlack1 channels (middle), and oocytes expressing eSlick channels (right) immediately after impaling the cell (start; black) and after 9 min of loading (end; gray). *B:* Whole cell currents recorded from the three types of oocytes mentioned above immediately after impaling the cell (start; red) and 9 min after (end; pink), with both microelectrodes filled with 2M KCl solution. *C:* Current amplitudes were normalized to the current recorded at “start”, and taken the log (base 2). Normalized current amplitudes from control cells (left), cells expressing eSlack1 (middle) and cells expressing eSlick (right)were plotted against time with 2M NaCl (black) or KCl (red) loaded to the cell. Measurements from 8 cells in each group were analyzed. Standard error was shown as gray or pink shades.

During a 9-min recording, the peak amplitude of both eSlack1 and eSlick at membrane potential of +20 mV was elevated with NaCl diffusing into the cell (Fig. 7A and C). To distinguish the role of Na^+^ and Cl^-^ in increasing current amplitude, we also used microelectrodes filled with 2M KCl and measured the peak amplitude of both eSlack1 and eSlick within 9 min. The currents of eSlack1 stayed constant with increased intracellular levels of KCl. In contrast, eSlick showed increased current magnitudes with KCl-filled electrodes (Fig. 7B and C). The peak current of control cells at +20 mV remained constant during 9-min loading of either 2M NaCl or KCl (Fig. 7). These results suggest that eSlack1 has an absolute requirement of Na^+^ to increase the channel’s open probability, whereas, eSlick channels are more sensitive to intracellular Cl^-^ levels.

### Transcription levels of eSlick increase with EOD frequency

*E. virescens* has considerable animal-to-animal variability in EOD frequency (200-500 Hz) [8] (Fig. 1E). Previous studies in a closely related species *Sternopygus macrurus* have shown that potassium channels in the EO are expressed in a gradient with EODf [21], this led us to examine whether the mRNA levels of *eSlack1, eSlack2* and *eSlick* genes in the EO vary across fish with different EOD frequencies. We extracted RNA from the EO of 10 fish with different EOD frequencies (192 Hz, 206 Hz, 229 Hz, 250 Hz, 300 Hz, 333 Hz, 350 Hz, 380 Hz, 395 Hz, 426 Hz) spanning most of the species’ natural range, and measured the transcription levels of the three K_Na_ channel genes with real-time PCR. We found that only the transcription level of eSlick was positively correlated with EOD frequency, whereas eSlack1 and eSlack2 were not transcribed in a gradient with EOD frequency (Fig. 8A and Fig. 9A-C)). We also divided the fish into two groups with high (≥ 300 Hz) and low (< 300 Hz) EOD frequencies, and compared the mean transcription level of the three K_Na_ channel genes between the two groups. Significant differences between fish with high and low EOD frequencies was only noted in the transcription level of eSlick (high frequency: n = 5, 3.52 (Mean) ± 0.58 (SEM); low frequency: n = 5, 1.94 (Mean) ± 0.28 (SEM),; Student’s t-test, *p* = 0.039), but not eSlack1 (high frequency:, n=5, 1.51(Mean) ± 0.40 (SEM); low frequency: n=5, 0.96 (Mean) ± 0.23 (SEM); Student’s t-test, *p* = 0.258) and eSlack2 (high frequency: n=5, 2.31(Mean) ± 0.31(SEM); low frequency: n=5, 1.83 (Mean) ± 0.95 (SEM); Student’s t-test, *p* = 0.647) (Fig. 8C).

**Fig. 8.**
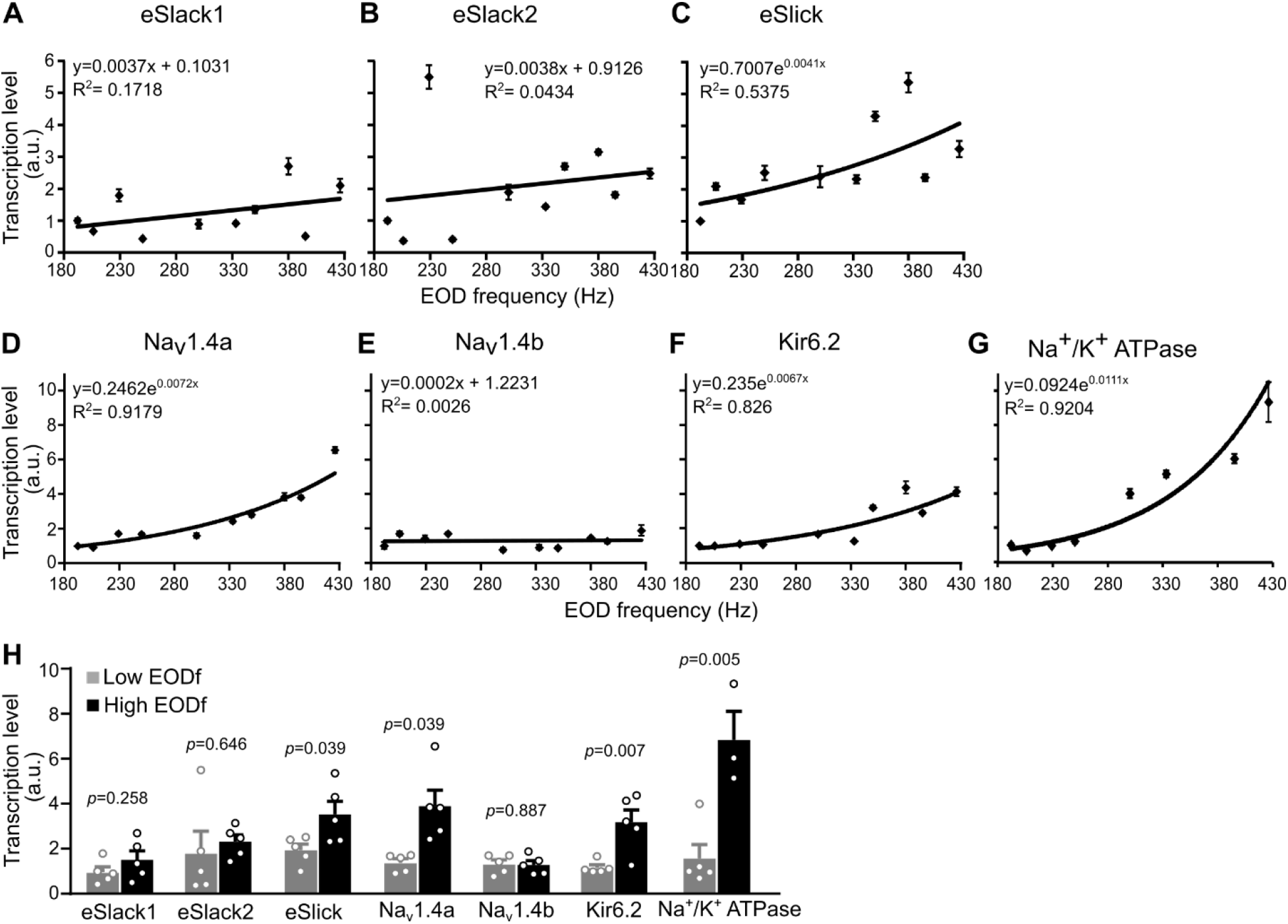
Real-time PCR quantification of ion channel genes in EOs from *E. virescens* with different EOD frequencies. *A-B:* The normalized transcription levels of target genes were plotted against EOD frequency. *A:* The transcription level of eSlack1 (A1) and eSlack2 (A2) in the EO do not correlate with EOD frequency. The transcription level of eSlick increases with EOD frequency (A3). *B:* The transcription levels of Nav1.4a (B1), Kir6.2 (B3) and Na^+^/K^+^ ATPase (B4) in EO from fish with different EOD frequencies can be fitted into an exponential curve. There is no correlation between the normalized amounts of Nav1.4b (B2) transcripts in EO and EOD frequency*. C:* Comparison of the mean transcription levels of genes between low (< 300 Hz) and high (> 300 Hz) frequency EOs. The average amounts of eSlick, Nav1.4a, Kir6.2 and Na^+^/K^+^ ATPase in high frequency EOs are higher than that in low frequency EOs. There is no significant difference in the mean transcription levels of eSlack1, eSlack2 and Nav1.4b between high and low frequency EOs..

**Fig. 9.**
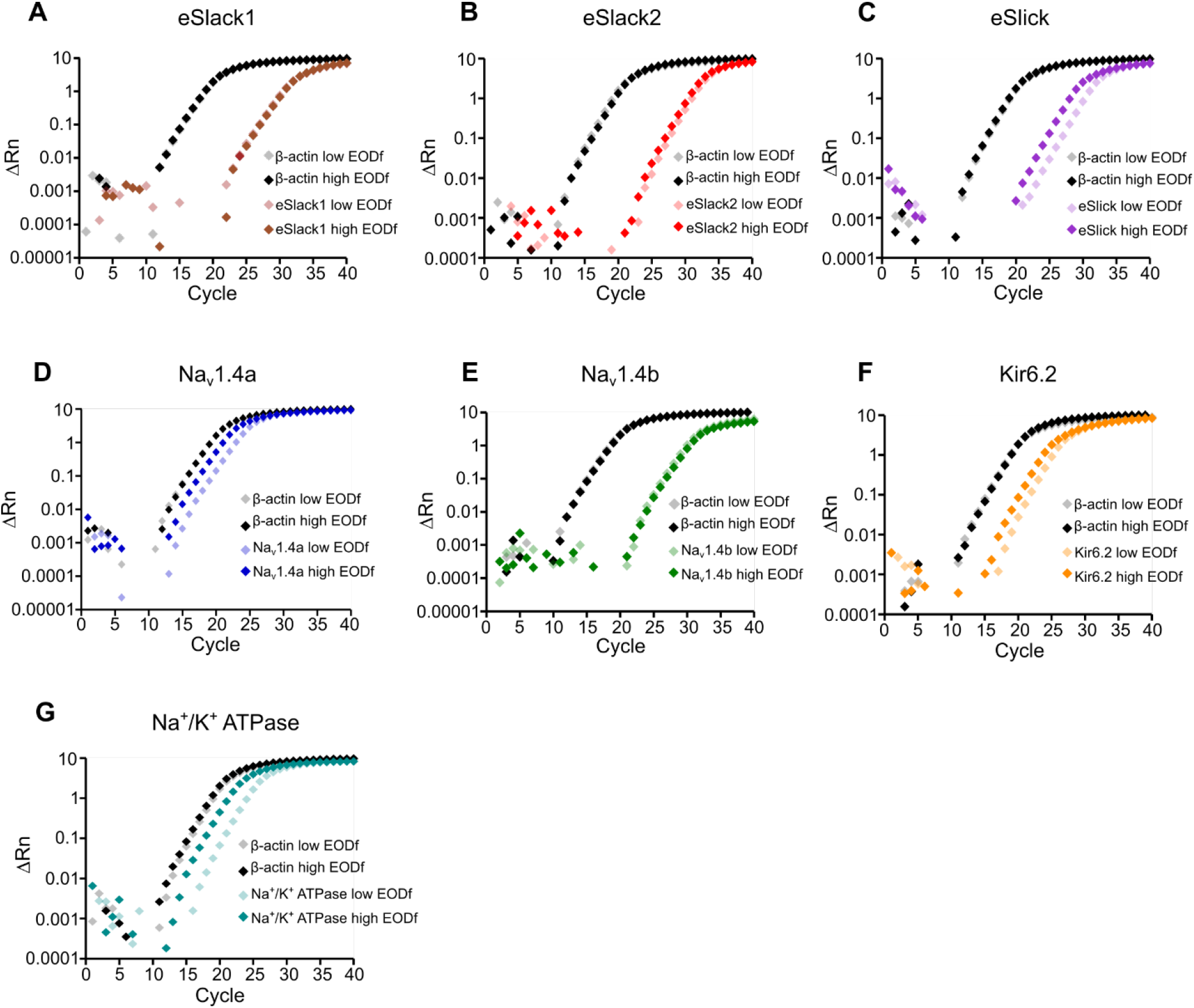
Amplification of the target genes and endogenous control β-actin from EO cDNA of a fish with low EODf and a fish with high EODf. The amplifications of β-actin, eSlack1 (A), eSlack2 (B), and Nav1.4b (E) from EO cDNAs of fish with high and low EODf look identical. eSlick (C), Na_v_1.4a (D), Kir6.2 (F), and Na^+^/K^+^ ATPase (G) started amplifying and reached the amplification plateau phase earlier when using EO cDNAs from a fish with high EODf than EO cDNAs from a fish with low EODf.

### Transcription levels of Na_v_1.4a, Kir6.2 and Na^+^/K^+^ ATPase increase with EOD frequency

In addition to the Na^+^-activated K^+^ current observed in electrocytes, whole cell recordings of endogenous currents in electrocytes also indicate the existence of an inwardly rectifying K^+^ (Kir) current and a voltage-gated Na^+^ (Na_v_) current [14]. The firing frequency of electrocytes is maintained by the coordination between ion channels involved in generating APs and the Na^+^/K^+^ ATPases, which are responsible for restoring the ionic gradients after each AP. The Na_v_ channels in the EO of *E. virescens* are encoded by a pair of duplicated genes, Na_v_1.4a and Na_v_1.4b, which are orthologs of the mammalian muscle-specific Na_v_1.4 gene [26]. The Kir channels are ATP sensitive potassium (K_ATP_) channels encoded by the KCNJ11 gene (unpublished data). In reverse transcription PCR, we noted that Na_v_1.4b and Kir6.2 are expressed in both muscle and EO, whereas, Na_v_1.4a and the α-subunit of Na^+^/K^+^ ATPases are dominantly expressed in the EO (data not shown). We reasoned that EOD frequency might also correlated with expression levels of these ion channels and Na^+^/K^+^ ATPases.

With real-time PCR, we measured the mRNA levels of these genes from the EOs of the same 10 fish used in the previous experiment. Results showed that the transcription levels of Na_v_1.4a, Kir6.2 and Na^+^/K^+^ ATPase increased exponentially with EOD frequency (Fig. 8B1, B3 and B4, and Fig. 9D, F and G). No correlation between the transcription level of Na_v_1.4b and EOD frequency was detected (Fig. 8B2 and Fig. 9E). When comparing the mean transcription level of genes between the two groups of fish with high and low EOD frequencies, significant difference was detected in Na_v_1.4a (high frequency: n=5, 3.89 (Mean) ± 0.72 (SEM); low frequency: n=5, 1.38 (Mean) ± 0.17 (SEM); Student’s t-test, *p* = 0.009), Kir6.2 (high frequency: n=5, 3.18 (Mean) ± 0.55 (SEM); low frequency: n=5, 1.16 (Mean) ± 0.13 (SEM); Student’s t-test, *p* = 0.007) and Na^+^/K^+^-ATPase (high frequency: n=3, 6.84 (Mean) ± 1.28 (SEM); low frequency: n=5, 1.57 (Mean) ± 0.61 (SEM); Student’s t-test, *p* = 0.005), but not Na_v_1.4b (high frequency: n=5, 1.28 (Mean) ± 0.19 (SEM); low frequency: n=5, 1.32 (Mean) ± 0.19 (SEM); Student’s t-test, *p* = 0.887) (Fig. 8C).

### Frequency-dependent scaling of multiple ionic conductances is necessary to maintain the integrity of simulated AP trains as EOD frequency increases

The frequency-dependent scaling of Nav1.4a, eSlick, and Kir6.2 mRNA levels led us to evaluate whether scaling the conductances associated with these mRNA transcripts is necessary to maintain electrocyte AP firing as AP frequency increases. In five model cells, we ran a series of simulations where AP frequencies from 150 Hz to 425 Hz in 25 Hz increments. (Fig. 10a, 10b). The ionic conductances of the five model cells were 1) all conductances fixed across AP frequencies, 2) only 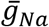 scaled exponentially with AP frequency, 3) both 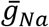 and 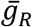 scaled with AP frequency, 4) both 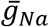 and 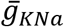 scaled with AP frequency and 5) a model where 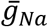, 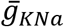, and 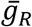 scaled with AP frequency. For model cells where ionic conductances were scaled, we scaled those conductances according to the best fit lines from the previous RNA expression data. In the first model where all conductances were fixed, AP amplitude decreased markedly at higher AP frequencies, a result of reduced membrane potential at peak, and incomplete repolarization before the subsequent AP (Fig. 10a,b,g-i). For the second and third models where only 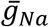, or where 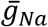, and 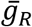 scaled with AP frequency, AP amplitudes were relatively stable through intermediate AP frequencies then declined precipitously at higher AP frequencies, largely because of near-failures to repolarize in the interspike interval (Fig. 10c,d,g-i). Allowing 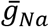 and 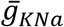 to scale with AP frequency resulted in highly stable AP amplitudes across all AP frequencies (Figure 10e,g-i) and this stability was improved when all three conductances 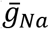, 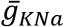, and 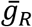) scaled with AP frequency (Fig. 10f-j). The improvements in AP consistency seen in the last two models was primarily a result of improved repolarization during the interspike interval (Fig. 10i), with the model producing the most consistent AP amplitudes being the one where all three conductances scaled with AP frequency (Figure 10j). These outcomes suggest that scaling these ionic conductances in a frequency-dependent manner consistent with our mRNA expression data is necessary to maintain AP waveform integrity as AP frequency increases.

**Fig. 10.**
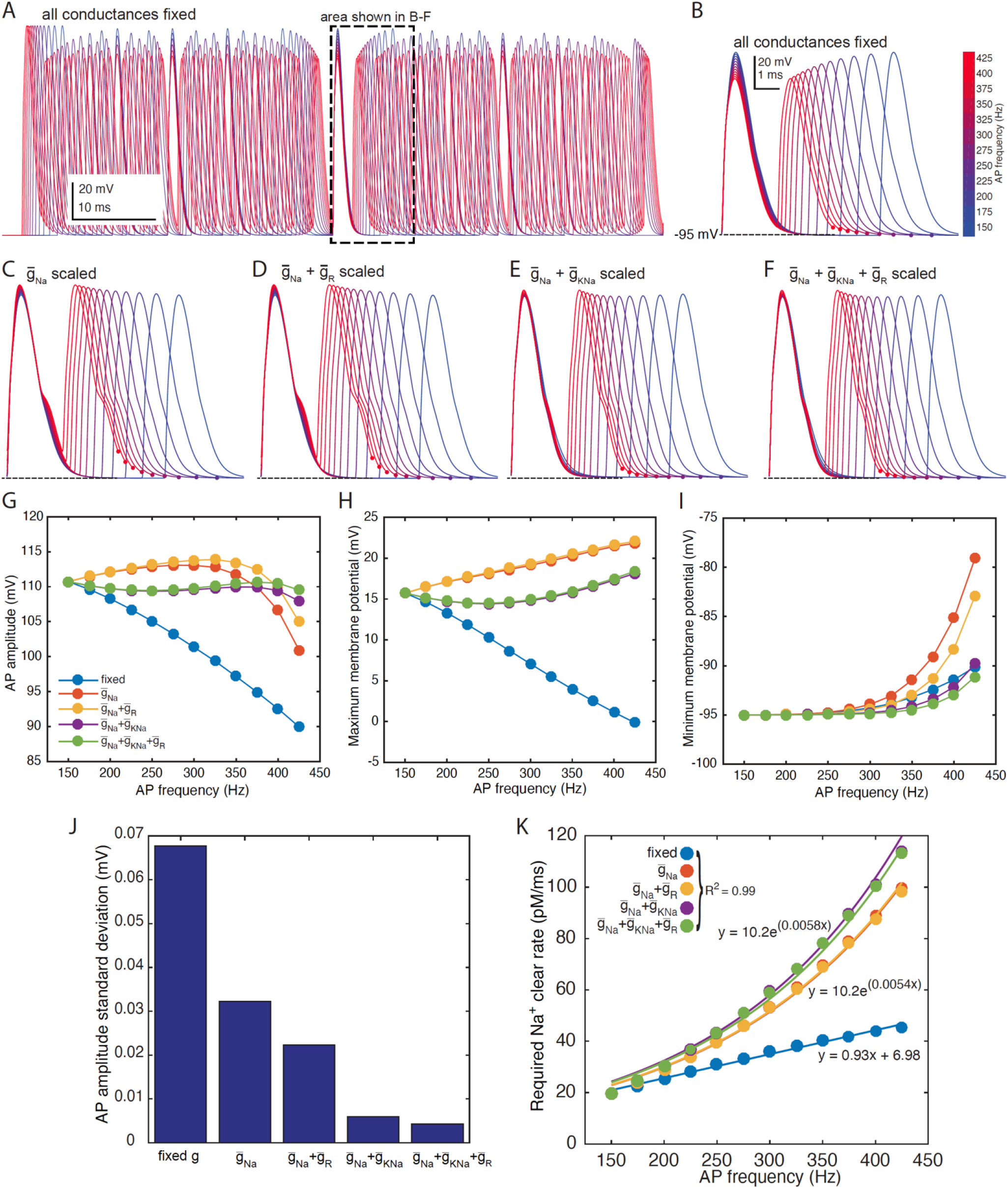
Computational simulations of electrocyte APs across AP frequencies in model electrocytes with and without scaled ionic conductances. (A) Superimposed trains of APs at frequencies from 150 Hz (blue) to 425 Hz (red) in 25 Hz steps, where all ionic conductances were held constant across AP frequencies The area within the dashed box is shown on expanded scale in *B* through *F*. (B) Two sequential action potentials within each of the AP trains shown in *A*. Firing frequency is indicated according to the color map at right from 150 Hz (blue) to 425 Hz (red). Small color-coded circles indicate where the next AP in the train initiated (subsequent APs not shown for clarity). All ionic conductances were constant across AP frequencies. At higher AP frequencies, peak membrane potential during the AP decreased and AP repolarization was increasingly incomplete during the interspike interval. (C) Sequential APs represented as in *B*, where 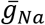 scaled exponentially with AP frequency according to the scaling equation 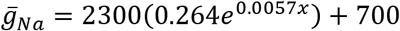. (D) Sequential APs where 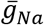 scaled with AP frequency as in *C*, and 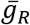 scaled with AP frequency according to the equation 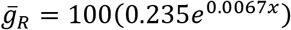. (E) Sequential APs where 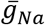 scaled with AP frequency as in *C* and 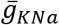 scaled with AP frequency according to the equation 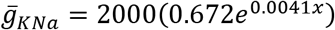. (F) Sequential APs where 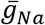 and 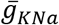 scaled with AP frequency as in *E* and 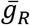 scaled with AP frequency as in *D*. (G) AP amplitude, measured peak-to-trough for five model cells across AP frequencies from 150 Hz to 425 Hz. Ionic conductances were fixed for all frequencies or various combinations of ionic conductances were varied in a frequency-dependent manner as indicated in the legend. AP amplitude decreased significantly at higher frequencies where conductances were fixed. Consistency of AP amplitudes improved as conductances were scaled with frequency, with the greatest consistency across frequencies produced by the model cell where 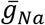, 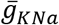, and 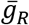 were all scaled with AP frequency. (H) Peak membrane potential during the AP for the same five model cells in *G*. Ionic conductances were fixed for all frequencies or various combinations of ionic conductances were varied in a frequency-dependent manner as indicated in the legend. Peak membrane potential decreased significantly at higher frequencies where conductances were fixed. Consistency of peak AP potential improved as conductances were scaled with frequency, with the greatest consistency across frequencies produced by the model cell where 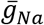, 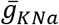, and 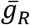 were all scaled with AP frequency. (I) Minimum membrane potential during the interspike interval for the same five model cells in *G*. Ionic conductances were fixed for all frequencies or various combinations of ionic conductances were varied in a frequency-dependent manner as indicated in the legend. Repolarization was most complete at higher frequencies for the model cell where 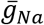, 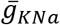, and 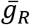 were all scaled with AP frequency. (J) Standard deviation of AP amplitude (as measured in *G*) across all AP frequencies for model cells where all ionic conductances were fixed, or scaled with frequency. The model cell with fixed conductances exhibited the largest variability in AP amplitude, while the model cell where 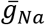, 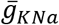, and 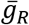 were all scaled with AP frequency showed the least variability in AP amplitude. (K) Required rate of Na^+^ extrusion to restore Na^+^ gradients (Na^+^ clear rate) for five model cells across AP frequencies from 150 Hz to 425 Hz. Na^+^ clear rate was computed as pM of Na^+^ entering the cell during the AP divided by the interspike interval. Ionic conductances were fixed for all frequencies or various combinations of ionic conductances were varied in a frequency-dependent manner as indicated in the legend. Filled circles are individual data points. Solid lines indicate least-squares regression fits. Goodness of fit, as measured by R^2^ was equal across all model cells. The Na^+^ clear rate increased in a linear fashion (equation shown in figure) for the model cell where all conductances were fixed. For cells where ionic conductances were varied with AP frequency, Na^+^ clear rates increased exponentially according to the equations shown in the figure. The exponential equations were indistinguishable within rounding for the cells where 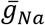, or 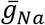 and 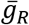, were varied as where the equations for cells where 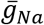 and 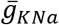, or where 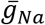, 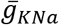, and 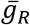, were scaled.

### Demand for Na^+^ transport increases exponentially as EOD frequency increases

Our simulations also suggest that an exponential scaling of Na^+^,K^+^-ATPase expression is necessary as EOD frequency increases. We calculated the required rate of Na^+^ extrusion to restore Na^+^ gradients (Na^+^ clear rate) for all five model cells across AP frequencies from 150 Hz to 425 Hz. Na^+^ clear rate was computed as the pumping rate necessary to return all Na^+^ that entered the cell during the AP to the extracellular space before the initiation of the next AP. The Na^+^ clear rate increased in a linear fashion (equation shown in figure) for the model cell where all conductances were fixed. For the other four cells in which ionic conductances varied with AP frequency, Na^+^ clear rates increased exponentially. The greatest increases in Na^+^ clear rate across AP frequencies occurred in the last two model cells where where 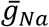 and 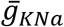, or 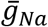, 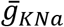, and 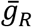 scaled with AP frequency, thereby producing the greatest consistency in AP amplitudes (Fig. 10j,k).

## DISCUSSION

*E. virescens* electrocytes face a challenge that is unique among excitable cells: maintaining unremitting firing rates of 200 to 600 Hz throughout the lifespan while producing microAmp-scale ionic currents during each AP. How can these cells maintain these extremely high firing rates and satisfy equally extreme demands for the rapid restoration of ion gradients between APs? Our findings here provide several key insights about the ionic mechanisms that make these feats possible as well as their metabolic consequences. We found that electrocytes fulfill the fast-spiking requirement by exponentially increasing expression levels of multiple ion channels, including a novel rapidly-activating K_Na_ channel isoform. Scaling expression levels of these channels to match firing rates is necessary for maintaining the integrity of the AP waveform as firing rates increase, but this comes at the cost of exponentially increasing demand for charge translocation by the Na^+^,K^+^-ATPase. Consistent with this conclusion we also found that Na^+^,K^+^-ATPase mRNA expression levels and electrocyte AP metabolic demand increase exponentially at higher firing rates.

We measured the mRNA levels of Na_v_1.4a, Na_v_1.4b, eSlack1, eSlack2, eSlick, Kir6.2 and Na^+^/K^+^ ATPase in EO from *E. virescens* with different EODf and found that transcription levels of Na_v_1.4a, eSlick, Kir6.2 and Na^+^/K^+^ ATPase are positively and exponentially correlated to EODf whereas levels of the other transcripts were not. Among the four genes that were correlated with EODf, Na_v_1.4a and Na^+^/K^+^ ATPase are predominantly expressed in EO and eSlick is expressed exclusively in EO, whereas Kir6.2 showed similar transcription levels in both muscle and EO, as was also the case for the genes that did not correlate with EODf (Na_v_1.4b, eSlack1, and eSlack2). Importantly, mRNA levels do not always predict its protein abundance, and we have not directly determined whether the abundance of corresponding proteins in electrocytes also correlates with EODf due to the lack of specific antibodies targeting most of the ion channels in electrocytes. However, our computational simulations strongly support the conclusion that expression of channel proteins in the cell membrane, in accordance with our observed mRNA expression levels, is required for the maintenance of AP waveform integrity across firing frequencies.

### Diverse roles of specific ionic conductances for fast-spiking in electrocytes

Given the faster activation kinetics for eSlick compared to eSlack1 and eSlack2, the correlation of eSlick with EODf is consistent with the briefer APs required as EODf increases. Higher expression of Na^+^/K^+^ ATPases at higher EODfs also is easily understood for higher frequency EODs, as increased rates of AP generation will require more rapid restoration of ionic gradients following each AP. Faster K^+^ channels are required to achieve fast-spiking. However, they will also increase the overlap between Na^+^ and K^+^ currents, further magnifying the energetic costs of AP generation. If minimizing energy consumption is one constraint governing the types of ion channels expressed in neurons [47], neurons with faster K^+^ channels should also express Na^+^ channels with a faster inactivation speed in order to reduce the overlap between Na^+^ and K^+^ currents. We have not yet determined the biophysical properties of Na_v_1.4a and Na_v_1.4b in this system, but Na_v_1.4a channels in weakly electric fish have numerous amino acid substitutions in regions associated with activation and inactivation [48].

The functional significance of scaling Kir6.2 expression levels with EODf is less clear. These channels are expressed at similar levels in EO and skeletal muscle unlike the other transcripts that scaled with EODf, and our computational simulations suggest that scaling Kir6.2 levels has only a minor role in maintaining integrity of the electrocyte AP waveform at high frequencies. One possibility is that Kir6.2 conductances are instead more important for responding to the metabolic state of the electrocyte. In neural, cardiac, and endocrine systems, Kir6.2 forms functional complexes with a sulphonylurea receptor (e.g. SUR1) belonging to the ATP-binding cassette (ABC) superfamily [49, 50]. These Kir6.2-SUR1 complexes in other systems are inhibited by physiological levels of ATP, increasing the channel’s open probability as intracellular concentrations of ATP fall and ultimately preventing AP initiation [51, 52].

The relationship between ATP availability and activity of the Kir6.2-SUR1 complex in electrocytes requires further study to determine its precise role in electrocyte function. The Na^+^ influx during each electrocyte AP exceeds 10 μA, requiring approximately 2 × 10^10^ ATP molecules for the Na^+^,K^+^ ATPases to restore the ionic gradients after each AP [11], approximately 100 times higher than that estimated for mammalian neurons [11, 53, 54]. The significant metabolic expense of a single AP, coupled with very high firing frequencies potentially exposes electrocytes to frequent changes in metabolic status. One appealing hypothesis is that these K_ATP_ channel complexes may form an endogenous protective system to stabilize the cell’s bioelectrical properties under metabolic stress as is the case in cardiac myocytes [52]. Under this scenario higher levels of Kir6.2 would be required in higher-frequency electrocytes to overcome the larger Na^+^ conductances necessary for higher frequency APs.

In an earlier study we characterized the electrophysiological properties of a K_Na_ conductance in *E. virescens* electrocytes without distinguishing multiple channel isoforms [14]. In the present study, we discovered the presence of three K_Na_ channel subunits expressed in electrocytes, eSlack1, eSlack2 and eSlick. Among these eSlack1 and eSlack2 closely resemble mammalian Slack channels, whereas eSlick appears to be a novel K_Na_ isoform with significant sequence divergence from mammalian Slick including the loss of a conserved ATP binding motif. Mammalian Slick channels contain an ATP binding motif in the C-terminal tail and can be directly inhibited by intracellular ATP [25]. Whether eSlick is regulated by ATP remains a question for future research, but the absence of a conserved ATP binding motif in eSlick suggests that it is not.

It is noteworthy that *E. virescens* electrocytes terminate their APs with K_Na_ channels whereas in all other electric fish where electrophysiology data are available and where EOD frequencies are much lower, electrocyte APs are terminated by Kv channels [14–17]. Fast-spiking vertebrate cortical neurons maintain high firing frequencies and reduce the metabolic cost of AP generation to near the theoretical minimum by tuning the kinetics of Na^+^ and K^+^ conductances to achieve rapid Na^+^ current inactivation and delayed onset of a rapidly-activating K^+^ conductance [55]. Similar mechanisms support high firing frequencies in brainstem auditory neurons [56]. In both cases, maintaining fast firing rates depends on voltage-gated K^+^ conductances from the Kv3 family of K^+^ channels. In computational simulations from earlier work, however, we found evidence that K_Na_ channels were necessary to support high-frequency APs in *E. virescens* electrocytes, while Kv3.1 channels were insufficient to maintain high firing frequencies [14].

Similar to mammalian K_Na_ channels, the opening of eSlack1 channels requires elevations of intracellular Na^+^, whereas, eSlick channels’ opening appears to be more dependent on intracellular levels of Cl^-^. These characteristics offer K_Na_ channels several advantages over Kv channels in cells with high firing rates. The accumulation of intracellular Na^+^ with high frequency stimulation may enhance activation of K_Na_ channels which could in turn serve as a negative feedback mechanism for the increased activity of Na^+^ channels. Additionally, K_Na_ channels may play a protective role against the inhibition of Na^+^/K^+^ ATPases under hypoxic conditions. The natural habitats of *E. virescens* include regions of low-oxygen waters and these fish are reported to have higher tolerance to hypoxic stress [57, 58]. Hypoxia-induced inhibition of Na^+^/K^+^ ATPases results in the increase of intracellular Na^+^ concentrations, which might enhance K_Na_ channel activity to increase the cell’s ability to react to metabolic stress arising from hypoxia or dietary energy shortfalls [59].

### Multiple K_Na_ channel subunits with distinctive roles

Functional potassium channels are tetramers of four subunits, and channels can consist of homotetramers or heterotetramers. Heterotetrameric K^+^ channels have functional properties that are typically intermediate between the properties of homomeric channels for each subunit. In the present study, eSlick currents were much faster than eSlack currents and therefore better suited for higher frequency electrocytes. The positive correlation of eSlick expression with EODf suggests two possibilities. One is that the ratio of eSlick homotetrameric channels to eSlack1 homotetrameric channels increases as EODf increases. A second possibility is that the ratio of eSlick to eSlack subunits within heterotetramers increases with EOD frequency. All three K_Na_ channels in *E. virescens* electrocytes are expressed on the cells’ anterior region, suggesting the possibility that they form heterotetrameric K_Na_ channels. Additionally, the failure of eSlack2 expressed alone to produce functional K_Na_ channels strongly suggests that this subunit occurs only within heterotetramers formed with eSlack1 and/or eSlick.

In mammalian systems, RNA alternative splicing gives rise to multiple Slack variant transcripts, Slack-A, Slack-B and Slack-M, which are regulated by alternative promoters and differ in the residues in their N-terminus [30]. The N-terminus of Slack-B is necessary for the trafficking of Slick subunits into the plasma membrane and they can form heterotetrameric K_Na_ channels[60]. eSlack2 and eSlick share similar N-terminal sequences with rat Slack-A and Slick. eSlack1 have an unique N-terminus, which is not identical to the N-terminus of any known mammalian Slack and Slick subunits. Heterogeneous expression of *E. virescens* K_Na_ channels in *X. laevis* oocytes showed that eSlack1 and eSlick can form functional homotetrameric K_Na_ channels and, although eSlack2 can be successfully trafficked into the plasma membrane, it could not conduct currents, which is likely due to the shorter C-terminal tail. Future biochemical studies with immunoprecipitation are necessary to examine the interactions among the three *E. virescens* K_Na_ channel subunits and the possibility to form heterotetrameric ion channels.

### Coregulation of ionic conductance densities as a general mechanism for fast-spiking cells

Matching ion channel expression levels to EODf occurs also in the closely related *Sternopygus macrucrus*, a weakly electric fish with lower range of EODfs (~50-200 Hz). Here, the transcription levels of two Kv channel subunits in EO (Kv1.1a and Kv1.2b) are correlated with EODf [61] and one Na_v_ subunit expressed in EO (Na_v_ 1.4b) is also correlated with EODf [62, 63]. Interestingly, in these cases the relationships between EODf and channel expression levels were linear, rather than exponential, suggesting the possibility that the much higher firing frequencies of *E. virescens* necessitated not only a shift to a very different molecular class of repolarizing K^+^ conductances, but also an escalation to exponential scaling of those conductances.

It is now well known that excitable cells modify the expression patterns of ionic conductances in order to maintain a particular functional state [64], but the cellular mechanisms that govern this process remain elusive [65–67]. A similarly intriguing question arises in the case of electrocytes in the present study. Do electrocytes respond to a given firing rate determined by the pacemaker nucleus through a cell-autonomous mechanism that appropriately tunes the expression levels of the necessary ion channels, or does some cell-extrinsic, perhaps endocrine, mechanism regulate both pacemaker firing rate and electrocyte ion channel expression? The contrasts between our present results and previous findings in *S. macrurus* highlight that whatever mechanisms govern the scaling of ionic conductances in electrocytes, these mechanisms are both surprisingly general (operating on different classes of ion channels and with different scaling rules across taxa) and also very specific (targeting only specific ionic conductance within the electrocyte, and assigning a specific scaling factor to each conductance). An interesting question raised by the present findings is whether ionic conductance densities in fast-spiking central neurons are actively tuned when their prevailing firing frequencies change during development or in an experience-dependent manner. Recent reports of developmental increases in metabolic efficiency for fast-spiking cortical neurons [55] suggest this may be the case.

Understanding the mechanisms regulating of ion channel expression levels as firing rates change is important not only in the context of electric sensory and communication signals in fish, but also for understanding how the performance of fast-spiking cells is tuned and energy-information tradeoffs are managed in other systems such as auditory processing networks, and neural systems more generally. This is especially true because in both cases a tradeoff between firing rates and metabolic cost appears to be a major force that shapes both the operational properties and the functional limits of these systems [68]. In some cases, fast spiking cells maintain the metabolic costs of AP generation near the theoretical minimum, even at very high firing rates [55], while in other cases the metabolic costs of AP generation increase exponentially as firing rates increase [11]. Comparative analyses of these processes across different taxa and different systems of excitable cells is an important step toward finding and understanding general and potentially convergent mechanisms that maintain high-frequency activity in fast-spiking cells and the rules that govern energy-information tradeoffs in bioelectric signaling systems.

## MATERIALS AND METHODS

### Animals and tissue harvesting

*E. virescens* (glass knifefish) were obtained from tropical fish importers (Gunpowder Aquatics, Wimauma, FL), and housed in tanks in a recirculating aquarium system at 28 ± 1°C with water conductivity 100-150 μS/cm. They were kept under 12 hour light: 12 hour dark cycle and fed *ad libitum* with live blackworms. The EO tissue was harvested by cutting off ~2cm section of the tail and removing the overlying skin. Skeletal muscle tissue was dissected from the hypaxial muscle after fish were euthanized by immersion in 2% eugenol solution in aquarium water.

All methods described were approved by the Institutional Animal Care and Use Committee of The University of Oklahoma and complied with the guidelines given in the Public Health Service Guide for the Care and Use of Laboratory Animals.

#### EOD frequency measurements

Fish were transferred to the EOD recording tank with two recording wires attached to the two opposite end walls and a ground wire located at one of the side walls. They were allowed to move freely while EODs were differentially amplified with a Cygnuys FLA-01 amplifier (Delaware Water Gap, PA) and EODf of the amplified signal was measured with a RadioShack digital multimeter set in frequency mode. To prevent the effects of temperature and water conductivity on the fish’s EOD frequency, the recording tank was placed in the aquarium room and filled with water from the same aquarium system where the fish was kept. Representative EOD waveforms recorded from two fish are shown in Fig.1E.

#### Molecular Biology

##### Reagents

The pSP64 Poly(A) vector, ImProm-II™ Reverse Transcription System and GoTaq^®^ DNA polymerase were purchased from Promega (Madison, WI). The RNA Clean & Concentrator™-5 was purchased from Zymo Research (Irvine, CA). The SMARTer^®^ RACE 5’/3’ Kit was purchased from Clontech Laboratories, Inc. (Mountain View, CA). All other molecular biology reagents were purchased from Thermo Fisher Scientific (Waltham, MA).

##### RNA and cDNA preparation

Tissues were homogenized using LabGEN 125 homogenizer (Cole-Parmer). Total RNA was extracted using TRIzol^®^ reagent and purified using RNA Clean & Concentrator™-5. Genomic DNA contamination was removed by incubating the total RNA with DNaseI at room temperature for at least 15 minutes. RNA quality was assessed by loading and running total RNA in a 1% agarose gel containing 0.5% bleach and SYBR^®^ Green II RNA Gel Stain [69]. One microgram of EO total RNA was reverse transcribed to cDNA with oligo(dT)15 primer using ImProm-II™ Reverse Transcription System. The concentration of RNA and cDNA was measured by Qubit fluorometer 2.0 (Thermo Fisher Scientific).

##### Cloning and sequencing of genes encoding *E. virescens* K_Na_ channels

cDNAs of interest were amplified by polymerase chain reaction (PCR) and 5’/3’ rapid amplification of cDNA ends (RACE). All PCR and RACE products were initially analyzed on 1% agarose gels stained with SYBR^®^ Safe DNA Gel Stain, purified and cloned into TOPO TA vector or pRACE vector. In each cloning, plasmids extracted from ten isolated individual colonies were sequenced by the Biology Core Molecular Lab at University of Oklahoma. Sequence results were used as a query to search the rat protein database using the online NCBI blastx tool to determine the molecular identity of amplified products [70].

##### eSlack1

A ~500-bp fragment of eSlack1 was amplified by nested PCRs using Platinum^®^ Taq High Fidelity DNA Polymerase with two pairs of degenerate primers designed against the highly conserved regions of published nucleotide sequences of *Slack* in other species (external primer pair: forward 5’-ARAGYTTYACCTWYGCYKCCTTY-3’ and reverse 5’-RYYTTYTSNBG YARMAGRTGGCA-3’; internal primer pair: forward 5’-AYAARAARTAYGGWGTRTGT HTG-3’ and reverse 5’ - GGMGAGCTSCCRATRTABGGMGA-3’). The thermocycler conditions were 94°C for 2 min, 30 cycles of 94°C for 30 sec, 55°C for 30 sec, and 68°C for 3 min, followed by a final extension step of 68°C for 10 min. The missing 5’end of eSlack1 cDNA was amplified by the following reactions: 1) A ~1-kb fragment was amplified by a 5’RACE reaction with a Slack degenerate primer (5’-GGMGAGCTSCCRATRTABGGMGA-3’) and an universal primer provided by the SMARTer^®^ RACE 5’/3’ Kit. 2) A ~500-bp fragment was amplified in a PCR with a forward degenerate primer (5’-GCCWTCBCAGCTSCTGGTGGT - 3’) targeting the signature sequence of the K^+^ selectivity filter and an eSlack1 specific reverse primer (5’-GCAAAGTCCTTCACCGCCCA-3’) designed from the partial cDNA fragment. 3) A 5’RACE PCR was carried out with an eSlack1 reverse primer (5’- TCACCTGACTGTCTGCCTCACATGGAC -3’) and the universal primer to amplify the start codon as well as the 5’untranslated region (UTR). The missing 3’end of eSlack1 including the stop codon and 3’UTR was amplified by a 3’RACE reaction with an eSlack1 forward primer (5’- CTACCCGTCCACA GCATCATCACTAGC-3’) and the universal primer. Sequences of the five eSlack1 fragments were aligned into a single contig (Geneious software, Biomatters Ltd, Auckland NZ). The full-length cDNA of eSlack1 was amplified with a forward primer (5’- ATATATAAGCTTTCTT TATTACCGAAGGTGTCCCTCCG-3’) derived from the 5’UTR and a reverse primer (5’- TATATATCTAGAGTTTCGGTTGATCAGGTCAGTTTAAAC-3’) derived from the 3’UTR. It was cut at the HindIII and XbaI sites introduced in the primers and cloned into pSP64 Poly(A) vector. Sequence of the insertion was confirmed to match the five overlapping PCR products. eSlack1 cDNA contains 3495 nucleotides.

##### eSlack2

The presence of eSlack2 was noted when sequencing the TOPO TA vectors inserted with the ~500-bp PCR product amplified with a forward degenerate primer targeting the K^+^ selectivity filter signature sequence and a reverse eSlack1 specific primer. Insert sequences from ten plasmids were aligned and assembled into two contigs using the Geneious software. The two contigs share 78.8% homology in nucleotide sequence, with residue differences dispersed along the entire region, and both of them share the highest homology with rat Slack (NCBI BLAST). We next performed 5’ and 3’ RACE reactions to amplify the missing 5’ and 3’ ends. A ~1.5-kb product was amplified in the 5’RACE reaction with a gene specific reverse primer (5’- GAGCTGACGCAGAGCACCACGTGTTT-3’), and a ~5 kb product was amplified in the 3’RACE reaction with a gene specific forward primer (5’- GCGTACCCACTCTGCCATGTT CAACC-3’). Sequences of the three PCR products were aligned to a single contig using Geneious. Full length eSlack2 cDNA was amplified with a forward primer (5’-ATATATGT CGACCTTCTTTACAATGATGGGAC-3’) targeting the 5’UTR and a reverse primer (5’- TATATAGGATCCCATTGGACAGTATGAATGAC-3’) targeting the 3’UTR. It was cut at the SalI and BamHI sites introduced to the primers and cloned into pSP64 Poly(A) vector. Sequence of the insert corresponded to the consensus sequence of the aligned contig. eSlack2 is composed of 3093 nucleotides.

##### eSlick

When amplifying the 3’end of eSlack2 with an eSlack2 specific forward primer (5’- TCTGGTGGTGGTGGACAAGGAGAGC-3’) and the universal primer, we detected a ~2.5-kb fragment. The RACE PCR product was cloned and sequenced as described. Nucleotide sequence was then blasted against rat protein database in NCBI, and shown to share the highest homology with rat Slick but not Slack. Then a 5’RACE PCR was performed to amplify the missing 5’end of the *Slick* transcript using a gene specific reverse primer (5’-ACGTCCTTATCCACAGAT CCTCCTCGG-3’). A ~4-kb DNA fragment was amplified, cloned and sequenced. Sequences of the two DNA fragments were aligned to a single contig containing potential start codon at the 5’ region and stop codon at the 3’ end. Full-length eSlick cDNA was amplified with a forward primer (5’- ATATATGTCGACTTTAGAGGAACGCATACTTAGC-3’) designed against the 5’UTR and a reverse primer (5’- TATATAGGATCCTAAGTAGTCAGATCAGTAGGGC-3’) designed against the 3’UTR. It was cut at the SalI and BamHI sites introduced in the primers and cloned into pSP64 Poly(A) vector. eSlick cDNA contains 3510 nucleotides.

##### Reverse transcription PCR analysis of gene expression in EO and muscle

To identify the expression patterns of target genes in EO and muscle, Reverse Transcription PCR was performed using GoTaq^®^ DNA polymerase with one microliter EO or muscle cDNA. Genes of interest and their specific primers are listed in Table 1. Thermocycling conditions included 95°C for 2 min, 35 cycles of 95°C for 30 sec, 55°C for 30 sec, and 72°C for 1min or 2 min (depending on the size of amplicons), and a final extension at 72°C for 5min. After gel electrophoresis, PCR products were visualized using the Safe Imager™ 2.0 (Thermo Fisher Scientific). Gel images were taken with the same exposure time.

**Table 1.**
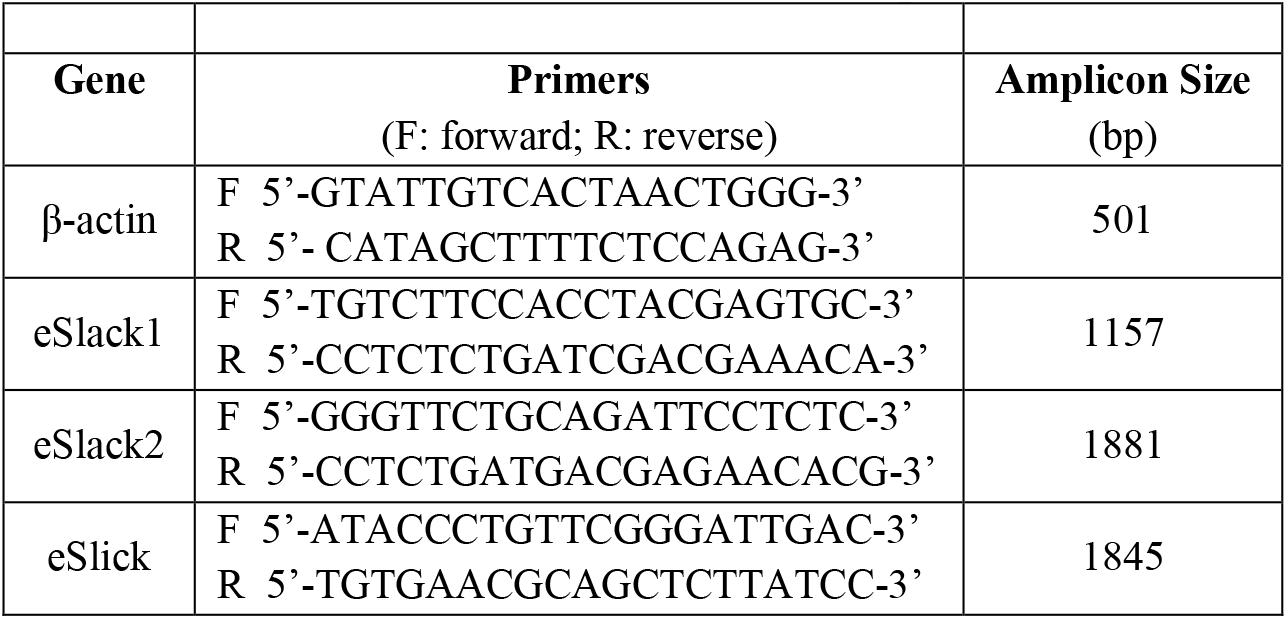
Primers used in reverse transcription PCR

##### Real-time PCR for mRNA quantitation in EO

One microgram of total RNA extracted from EOs from 11 adult *E. virescens* with EOD frequencies (192Hz, 202Hz, 206Hz, 229Hz, 250Hz, 300Hz, 333Hz, 350Hz, 380Hz, 395Hz, 426Hz) spanning the species’ natural range was reverse transcribed to cDNA with oligo(dT)15 primer using ImProm-II™ Reverse Transcription System. cDNA was diluted to 20 ng/μl. Gene specific primers were designed using the GenScript online software to control the primer length ~20 bases, melting temperature (Tm) in the range of 58-60°C, and amplicon size ~100 bp (Table 2). Each reaction contained 100 ng cDNA, 25ul 2× Power SYBR^®^ Green Master Mix, 200 nM of forward and reverse primer, and nuclease-free H_2_O to reach a total volume of 50 μl. Experiments were run in an Applied Biosystems 7500 Real-time PCR system using the default run method for Power SYBR^®^ Green cDNA two step kit: hold at 95°C for 10 min and 40 cycles amplification (denature at 95°C for 15 sec, and anneal/extend at 60°C for 1min). Each sample has three technical replicates. The specificity of primers was assessed by both melt curve analysis and gel electrophoresis of qPCR product. The expression level of all target genes were normalized to the endogenous control β-actin. EO cDNA from fish with the lowest EOD frequency (192 Hz) was used as the calibrator sample. Reactions without cDNA template were performed as negative controls. All negative controls showed no amplification or amplification starting more than eight cycles later than the reactions with cDNA template. In each experimental run, the standard curve was generated using 500, 100, 20, 4, 0.8 ng EO cDNA from a fish with 202Hz EOD frequency. All target genes and β-actin had standard curves with R^2^>0.97. The slopes of standard curves were used to estimate the amplification efficiencies, which were in the range between 95% and 105%. Data were analyzed using Applied Biosystems 7500/7500 Fast software. Standard deviations were calculated by following the Applied Biosystems guide to perform relative quantification of gene expression using relative standard curve method.

**Table 2.**
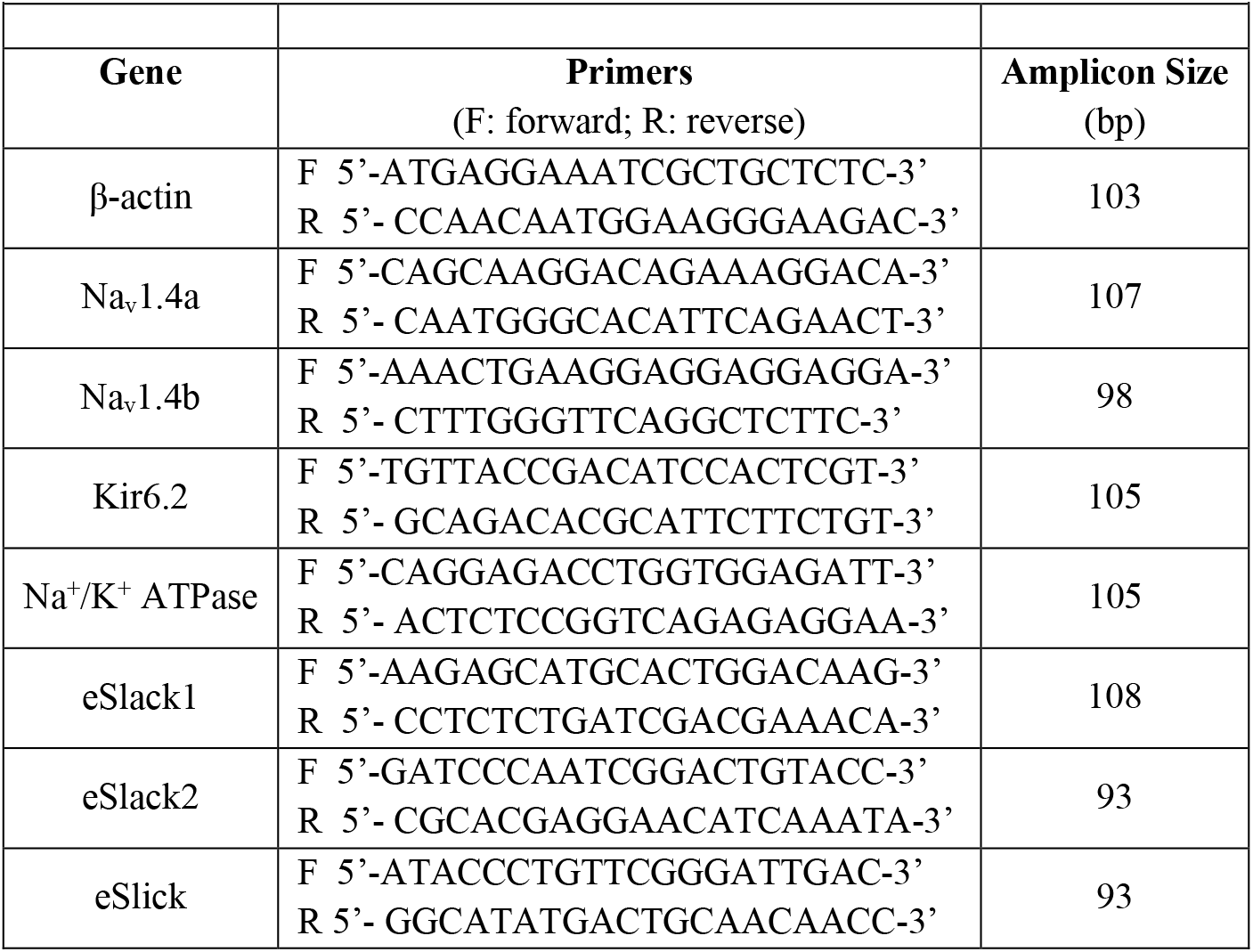
Primers used in real-time PCR

#### Gene phylogeny analysis

*E. virescens Slack1, Slack2*, and *Slick* cDNA sequences were translated and aligned with protein sequences of the SLO family channels in nematode, zebrafish, mouse, rat and human using ClustalW. Then the phylogenetic relationship was analyzed using the Geneious software (version 7.1.7). The consensus tree was obtained by using neighbor-joining method, Jukes Cantor amino acid substitution model and resampled 1000 times with Bootstrapping method. Human voltage gated K^+^ channel subfamily A member 1 (hKv1.1) was included as the outgroup. Channels included in the phylogenetic analysis are *Caenorhabditis elegans* Slo1channel (NCBI accession number: Q95V25); *Danio rerio* Slo1 channel (NP_001139072); *Mus musculus* Slo1 channel (NP_001240287); *Rattus norvegicus* Slo1 channel (NP_114016); *Homo sapiens* Slo1 channel (AAI44497); *Caenorhabditis elegans* Slo2 channel (AAD51350); *Danio rerio* Slack channel (XP_009293403); *Danio rerio* Slick channel (XP_017214614); *Mus musculus* Slack channel (NP_780671); *Mus musculus* Slick channel (NP_001074496); *Rattus norvegicus* Slack channel (NP_068625); *Rattus norvegicus* Slick channel (NP_942057); *Homo sapiens* Slack channel (NP_065873); *Homo sapiens* Slick channel (NP_940905); *Mus musculus* Slo3 channel (O54982); *Homo sapiens* Slo3 channel (NP_001027006); *Homo sapiens* Kv1.1 channel (NP_000208).

#### Expression of recombinant K_Na_ channels in electrocytes

We constructed recombinant eSlack1, eSlack2 and eSlick channels tagged with the red fluorescent protein (mCherry) at their N-terminus. mCherry was PCR amplified from u-mCherry (a gift from Scott Gradia; Addgene plasmid # 29769). The polylinker sequences between ion channels and mCherry are GGSGGGSGGSGS for eSlack1/ eSlick, and GGSGGGSG for eSlack2 [40, 41]. mCherry-eSlack1, mCherry-eSlack2 and mCherry-eSlick was assembled and cloned into pOX vector using the NEBuilder^®^ HiFi DNA Assembly Master Mix (New England Biolabs^®^ Inc.), then subcloned into pmaxCloning™ vector (Lonza). Prior to EO injection, the fish were anesthetized by exposing them to 0.01% clove oil until losing equilibrium but still maintaining opercular beating (< 2 min total). A single 25 μl bolus of 5 μg/μl plasmid in 150 mM KCl was injected into the fish’s EO in the tail using a microliter syringe. The injected fish was transferred to a bucket containing aerated water from its home tank and monitored for recovery, then transferred back to its home tank after full recovery. The expression of mCherry tagged ion channels was examined at the 10^th^ day after injection using epifluorescence and confocal microscopy.

#### Image acquisition

To examine the expression of mCherry-eSlack1, mCherry-eSlack2 and mCherry-eSlick in electrocytes, we harvested the EO using the same procedure as described earlier [37]. Live electrocytes were first examined on a Zeiss Apotome.2 microscope with a X5/0.16NA dry objective and processed by Zeiss AxioVision Rel.4.8.2. Structured illumination was used to create optical sections of the sample. Then we used LeicaTCS SP8 laser scanning confocal microscope with a X25/0.95NA dipping objective to acquire high resolution images. mCherry was excited by a 561-nm laser line and autofluorescence of electrocyte was excited by a 488-nm laser line [37]. The images were acquired as serial sections and processed by the software Leica Application Suite advanced Fluorescence (LAS AF) 3.3.0.10134. Electrocytes without expressing mcherry tagged eSlack/Slick subunits were used as control and imaged under the same settings.

*Xenopus laevis* oocytes expressing mCherry tagged K_Na_ subunits were incubated in ND96 saline and imaged using LeicaTCS SP8 laser scanning confocal microscope with a X10/ 0.3NA dry objective. Brightness and contrast of all images were adjusted using ImageJ for 64-bit Windows (version 1.51s; National Institute of Health).

#### Electrophysiology

eSlack/Slick cDNA was subcloned into pOX vector (a generous gift of Dr. Lawrence B. Salkoff, Washington University, St. Louis, USA). In vitro transcribed RNA (cRNA) was prepared using the mMESSAGE mMACHINE™ T3 Transcription Kit (Thermo Fisher Scientific). We used an Agilent 2100 Bioanalyzer to examine the quality and concentration of cRNA. Defolliculated *X. laevis* oocytes in stage VI were obtained from Ecocyte Bioscience (Austin, TX) and incubated in modified Barth’s saline containing the following in mM: NaCl 88, KCl 1, NaHCO_3_ 2.4, MgSO_4_ 0.82, Ca(NO_3_)_2_ ·4H_2_O 0.33, CaCl_2_ ·2H_2_O 0.41, HEPES 5, CH3COCOONa 2.5, and 50 μg/ml gentamycin at pH 7.5. Oocytes were injected with 46 nl of nuclease-free water containing ~80ng of cRNA and analyzed 4 to 5 days post injection.

Whole cell currents from oocytes were recorded using a standard two electrode configuration [14], with an Axoclamp 900 amplifier controlled by a Digidata 1440 interface and pCLAMP10 software (Molecular Devices, Sunnyvale, CA). Data were sampled at 100 kHz and filtered at 10 kHz. Electrodes were pulled from 1.2 mm o.d. thin-wall borosilicate glass tubing, filled with 2M NaCl or KCl and had resistances of 0.5-1.2 MΩ. Oocytes were incubated in ND96 saline (in mM: 96 NaCl, 2 KCl, 1 MgCl_2_, 1.8 CaCl_2_ ·2H_2_O and 5 HEPES, pH to 7.5). To measure channel activation, oocytes were held at −90 mV, then depolarized by 400 ms voltage steps ranging from −90 mV to +80 mV in 10 mV increments every 5 s. In some experiments, cells were depolarized by a 500 ms pulse to +20 mV from a holding potential of −90 mV every 10s to examine the effects of NaCl and KCl on the amplitude of whole-cell currents. The activation τ for eSlack and eSlick currents was estimated using the Clampfit fitting functions. Current traces from the start point to the peak point just before the plateau stage were fitted to a standard single-term exponential growth function. The time required to reach 62.8% of the final value was calculated as the activation τ.

#### Computational Simulations

We modeled *E. virescens* electrocyte APs with a simplified version of our earlier electrocyte simulations [37]. Briefly, the electrocyte was simulated with the Hodgkin-Huxley formalism as a three-compartment cell with an active posterior compartment, a passive central compartment, and an active anterior compartment. Simulated cholinergic synaptic current was applied only to the posterior compartment and the frequency of the synaptic inputs was varied to elicit trains of simulated APs with frequencies of 150Hz to 425Hz in 25 Hz increments.

The capacitances for the posterior, central, and anterior compartments were 48.0 nF, 18 nF, and 18 nF, respectively, based on surface area measurements from high-resolution confocal 3D reconstructions of single electrocytes. Differential equations integrated via Euler’s method were coded in Matlab (Mathworks, Inc. Natick MA) with integration time steps of 5 × 10^-8^ sec. The passive central compartment’s current balance equation included only passive leak (IL) fixed at 5 μS, and coupling to the two adjoining active compartments as in Equation 1

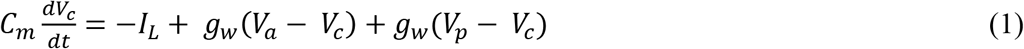

where *g_w_* is the coupling conductance, fixed at 3 μS. The current balance equation for the posterior compartment was

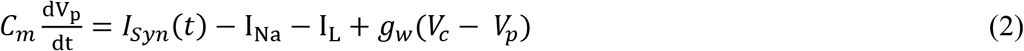

and the current balance equation for the anterior compartment was

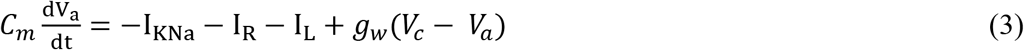

where ISyn represents synaptic current, I_Na_ is the voltage-gated Na^+^ current, IK_Na_ is the Na^+^-activated K^+^ current, I_R_ is the inward rectifier K^+^ current, and gw is the coupling current to the adjacent compartment. For all three compartments the leak current, IL, was given by Equation 4, where 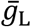 was 10 μS, 5 μS, and 10 μS for the posterior, central, and anterior compartments, respectively.

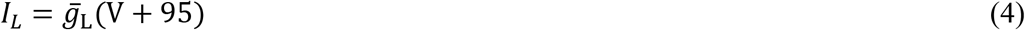

The posterior-compartment synaptic current, ISyn, was given by Equation 5

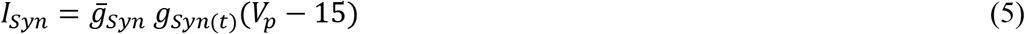

with 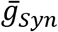 fixed at 600 μs for all models and where the time series gSyn(t) was a series of alpha waveforms generated using the discrete time equation [71]:

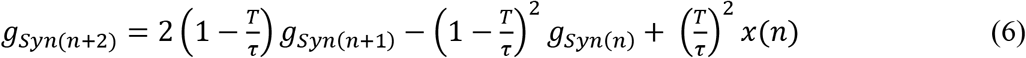

For this equation *T* is the integration time step and τ is the time constant. The binary series *x*(*n*) specified the onset times of the synaptic inputs, and the resulting time-series *g_Syn(n)_* was normalized such that 0 ≤ *g_Syn(n)_* ≤ 1.

The voltage-dependent currents I_Na_, IK_Na_, and I_R_ were given by Equations 7 – 9:

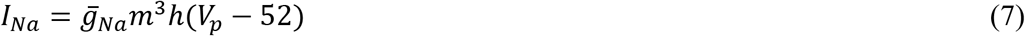

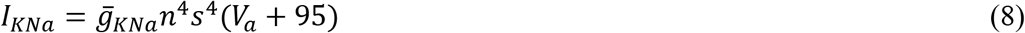

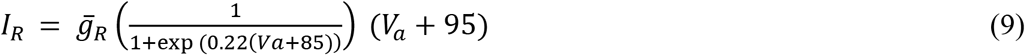

The baseline values of 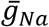, 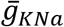, and 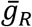, were 2300 μS, 2000 μS, and 125 μS, respectively.

The gating variables *m*, *h*, and *n* in Equations 7 and 8 evolved in a voltage-dependent manner according to Equation 10 where *V* is the membrane potential of the appropriate compartment (*V_p_ or V_a_*) and *j* = *m*, *h*, or *n*

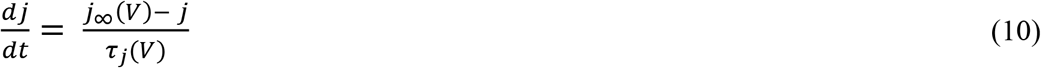

The voltage-dependent values of *j_∞_* in Equation 10 were determined according to Equations 11–13 for *j* = *m*, *h, or n, respectively*:

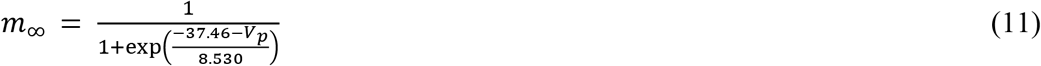

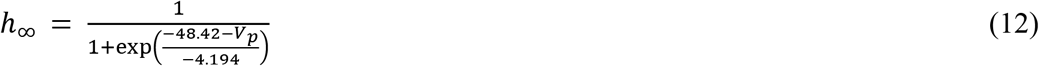

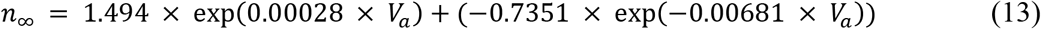

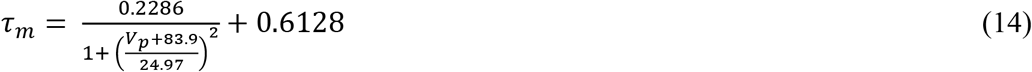

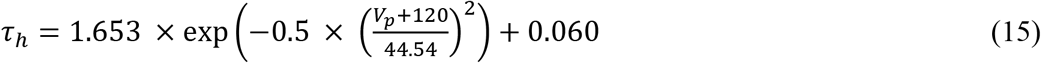

and τ_*j*_ was given by Equations 14, 15, and 16 for *j* = *m, h*, or *n*, respectively.

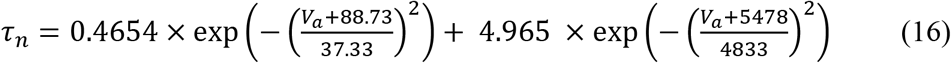

All parameter values in Equation 9 and Equations 11–16 were determined by least-squares best fits to experimental data for IK_Na_ from the present study for *j* = *n* and were determined by least-squares best fits to previous experimental recordings of I_Na_ in *E. virescens* electrocytes [14] for *j* = *m* and *h*. We previously modeled the Na^+^-dependence of gK_Na_ with the gating variable, *s*, which is determined by the Na^+^ concentration in the bulk cytoplasm in the anterior compartment [37]. In those simulations, however, there were no significant changers in Na^+^ concentration. We therefore did not model changes in Na^+^ concentrations in the present model and the Na+ gating variable *s* was therefore fixed at 0.7895, in accordance with the fixed anterior compartment Na^+^ concentration of 15 mM.

The values of 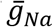, 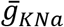, and 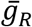, were scaled according to AP frequency in some models, and held constant in other models. Values of these parameters were given by Equations 17–19, where *x* denotes AP frequency:

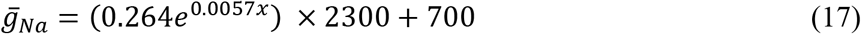

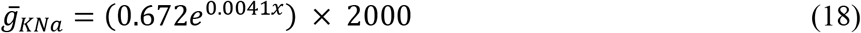

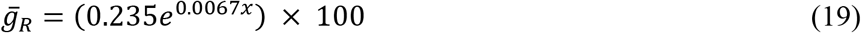

The base and exponential parameters in Equations 17–19 are based on RNA expression data from the present study. The remaining constants in each equation were selected to produce an AP train where the AP duration was one-half of the interspike interval at an AP frequency where *x* = 150 Hz. In models where values of 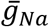, 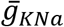, and/or 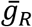, were held constant, the value of x was fixed at 150 regardless of AP frequency.

To determine the required activity of the Na,K-ATPase to restore ionic gradients between APs, we calculated total Na^+^ entry via the voltage-gated Na+ conductance during each AP as moles of Na^+^ by multiplying the integrated Na^+^ current (in nA*ms) by 10^-12^ to yield Coulombs of charge, then dividing by the elementary charge on a monovalent cation, *e*, to yield the number of Na^+^ ions, then dividing by Avogadro’s constant, *L*, to yield moles of Na^+^.

## ACKNOWLEDGEMENTS

The authors declare no conflict of interest and no competing financial interests. Financial support and equipment were provided by NSF grants IOS1350753, and IOS 1644965 (M.R.M.). This research was also supported in part by a grant from the Research Council of the University of Oklahoma Norman Campus. We thank in particular Lawrence Salkoff for the gift of pOX vector, David McCauley for valuable suggestions on experimental design, and Tingting Gu for imaging assistance. Thanks to Austin McCauley and Shannon Wiser for fish care, Tian Yuan and Mehrnoush Nourbakhsh for help in molecular cloning, Rosemary K_Na_pp for use of her cryostat, and J.P. Masly for use of his microscope.

